# Exercise alters cortico-basal ganglia network functional connectivity: A mesoscopic level analysis informed by anatomic parcellation defined in the mouse brain connectome

**DOI:** 10.1101/2023.02.23.529814

**Authors:** Zhuo Wang, Erin K. Donahue, Yumei Guo, Michael Renteln, Giselle M. Petzinger, Michael W. Jakowec, Daniel P. Holschneider

## Abstract

The basal ganglia are important modulators of the cognitive and motor benefits of exercise. However, the neural networks underlying these benefits remain poorly understood. Our study systematically analyzed exercise-associated changes in functional connectivity in the cortico-basal ganglia-thalamic network during the performance of a new motor task, with regions-of-interest defined based on mesoscopic domains recently defined in the mouse brain structural connectome. Mice were trained on a motorized treadmill for six weeks or remained sedentary (control), thereafter undergoing [^14^C]-2-deoxyglucose metabolic brain mapping during wheel walking. Regional cerebral glucose uptake (rCGU) was analyzed in 3-dimensional brains reconstructed from autoradiographic brain sections using statistical parametric mapping. Functional connectivity was assessed by inter-regional correlation of rCGU. Compared to controls, exercised animals showed broad decreases in rCGU in motor areas, but increases in limbic areas, as well as the visual and association cortices. In addition, exercised animals showed (i) increased positive connectivity within and between the motor cortex and caudoputamen (CP), (ii) newly emerged negative connectivity of the substantia nigra pars reticulata with the globus pallidus externus, and CP, and (iii) reduced functional connectivity of the prefrontal cortex (PFC). Increased functional connectivity in the motor circuit in the absence of increases in rCGU strongly suggests greater network efficiency, which is also supported by the reduced involvement of PFC-mediated cognitive control during the performance of a new motor task. Our study delineates exercise-associated changes in functional circuitry at the subregional level and provides a framework for understanding the effects of exercise on new motor learning.

## 1. INTRODUCTION

It is well documented that exercise improves brain cognitive, motor, and affective functions in health and disease, and has preventive and restorative benefits in neuropsychiatric conditions, as well as in age-associated functional decline (Cotman and Berchtold, 2002; Hillman et al., 2008; Petzinger et al., 2013; Gomes-Osman et al., 2018; Ludyga et al., 2020; Dauwan et al., 2021). At the molecular and cellular level, exercise effects are mediated by brain-derived neurotrophic factor (BDNF) and other signaling molecules, and expressed in multiple forms of brain plasticity, including neurogenesis, synaptogenesis, angiogenesis, improved mitochondrial function, altered neuroexcitability, improved or preserved white matter integrity, and enhanced neuroplasticity (Hillman et al., 2008; Nicolini et al., 2021). While these changes are believed to be beneficial to brain functions in general, how they lead to behavioral improvement remains incompletely understood. Neuroimaging investigation can offer insight of exercise effects by examining changes in functional connectivity on neural networks, thereby bridging the gap between microscopic neural substrates and behavioral outcomes (Won et al., 2021; Moore et al., 2022).

It is hypothesized that exercise, which involves motor and often cognitive tasks, recruits the cortico-basal ganglia-thalamic (CBT) network to bring about activity-dependent neuroplasticity at local and distant brain sites. Animal research has shown exercise-related metabolic, perfusion and molecular effects in individual regions of the CBT, in particular the caudoputamen (CP), motor cortex, substantia nigra, and thalamus. Yet, only a few studies have examined functional connectivity changes in the CBT network following exercise. We previously reported that in rats with bilateral 6-hydroxydopamine lesion to the CP, exercise partially reinstated cortical sensorimotor functional connectivity lost following dopaminergic deafferentation (Peng et al., 2014) and strengthened connectivity in the CBT and cerebellar-thalamocortical circuits (Wang et al., 2015). Ji et al. (2017) reported an exercise-associated increase in resting-state functional connectivity between putamen and thalamus in human subjects. Increased resting-state functional connectivity of the motor cortex has been reported in normal volunteers following several minutes (McNamara et al., 2007; Sun et al., 2007) or 4 weeks of motor training (Ma et al., 2010). Exercise-associated reduction in resting-state functional connectivity of the basal ganglia has also been reported (Magon et al., 2016; Tao et al., 2017). These studies typically report functional connectivity changes of large areas such as the whole putamen, masking subregional (mesoscopic level) heterogeneity in network structure and function. To our best knowledge, there has not been a systematic analysis of exercise-associated changes in functional connectivity at the mesoscopic level over the CBT network.

Recent development in the mouse brain connectome has brought unprecedented, detailed information on the structural organization of the CBT network. In particular, newly identified, domains at the mesoscopic level have been defined for key structures of the basal ganglia based on patterns of axonal projections. Dong and coworkers have subdivided CP into 29 domains based on the structural cortico-striatal projectome (Hintiryan et al., 2016), and further subdivided the globus pallidus externus (GPe) into 36 and substantia nigra pars reticulata (SNr) into 14 domains based on projections from the CP (Foster et al., 2021). This connectomic information creates a framework for systematic functional connectivity analysis of the basal ganglia. The current study applied the classic [^14^C]-2-deoxyglucose (2DG) uptake autoradiographic method of cerebral metabolic mapping to examine functional reorganization in the CBT network in response to chronic exercise. The well-established 2DG method is particularly suitable for high-resolution mapping in awake, freely-moving animals. We applied the newly identified domain definitions of Dong and coworkers in a region-of-interest approach to investigate exercise-associated changes in functional connectivity of the network. Following 6 weeks of exercise training on a motorized horizontal treadmill, glucose uptake was mapped in animals performing a novel, wheel-walking task. There is evidence that chronic exercise promotes motor learning capacity (Li and Spitzer, 2020), with associated changes in functional brain activation correlating with changes in aerobic fitness (Duchesne et al., 2016). Prior work has not examined correlation between aerobic fitness and functional connectivity in the CBT network, which is implicated in new motor learning (Dayan and Cohen, 2011). Our findings provide new insight into how exercise differentially alters functional interactions, both within individual structures and across the CBT network to improve learning capacity. Functional neuroimaging research that harnesses state-of-the-art anatomical connectomic information can bridge the current knowledge gap in understanding exercise effects between the microscopic and behavioral levels.

## 2. MATERIALS AND METHODS

### 2.1. Animals

Male C57BL/6J mice were purchased from Jackson Laboratory (Bar Harbor, Maine, USA) and housed in groups of 4-5 per cage on direct woodchip bedding at the University of Southern California vivarium. Animals had *ad libitum* access to laboratory rodent chow and water and were maintained on a 12-hr light/12-hr dark cycle (lights on at 0700 and off at 1900 hours). All experimental procedures involving animals were approved by the Institutional Animal Care and Use Committee at the University of Southern California (Protocol # 21044) and carried out in compliance with the National Institutes of Health Guide for the Care and Use of Laboratory Animals, 8th Edition, 2011.

### 2.2. Overview (Fig. 1)

A total of 20 mice were randomized into two groups (*n* = 10/group, aged 4 -6 months by the end of experiment): control (sedentary) and exercise. Animals received 6 weeks of exercise training on a motorized treadmill or sedentary treatment, followed by [^14^C]-2DG cerebral metabolic mapping while the animals walked in a wheel, a new motor task. The brains were cryosectioned into coronal slices, which were subsequently exposed to films for autoradiography. Digitized images of brain slices were used to reconstruct three-dimensional (3D) brains. These 3D brains were preprocessed using the Statistical Parametric Mapping (SPM) software, followed by statistical tests for between-group differences in regional cerebral glucose uptake (rCGU). For functional connectivity analysis, a total of 176 regions of interest (ROIs) were defined to represent mesoscopic-level domains of the CP, SNr, GPe, and to represent selected cortical and thalamic structures associated with motor and cognitive functions. Pairwise inter-regional correlation matrices were calculated for both groups to assess functional connectivity. Network organization was further analyzed using graph theory analytic tools.

### 2.3. Treadmill exercise

Mice were exercised on motorized treadmills (EXER-6, Columbus Instruments, Columbus, OH, USA) for 1 hr/day, 5 days/week over 6 weeks, as previously described (Lundquist et al., 2019) with slight modifications. The exercise protocol included a 20-min warm-up phase when a base speed of 5 m/min was ramped up incrementally every 5 min to a top speed, a 10-min running at the top speed, a 5 min walk break at 5 m/min, another 10-min running at the top speed, and a final 20-min cool-down phase when speed was ramped down every 5 min to a final speed of 5 m/min. During the first several days of exercise, top speed was incrementally increased to allow the mice to adapt to running at higher speed. The final top speed of 19 m/min was introduced on day 8 of exercise. Control sedentary mice were kept in home cages placed on a plastic barrier overlying the motorized treadmills for the same duration (1 hr/day, 5 days/week, 6 weeks) so that they were subjected to similar vibrations and auditory stimulation.

### 2.4. Autoradiographic glucose metabolic mapping during wheel walking

The autoradiographic 2DG uptake method is a well-established approach to functional brain mapping based on a tight coupling between neural activity and metabolism. It is particularly suitable in awake, freely-moving animals, and hence can be applied to exploration of network connectivity in the behaving animal. The protocol is as previously described with modifications (Sokoloff et al., 1977; Holschneider et al., 2019; Needham et al., 2022). For 2 days prior to the day of 2DG mapping, mice were individually familiarized to walk in a closed wheel for 10 min/day at a modest speed of 2 m/min (3.3 cm/s) on a motorized wheel bed (Model 80805A, Lafayette Instrument, Lafayette, IN, USA) — a new motor task that can be learned by all animals at this low speed. Using the wheel-walking task avoided the confound of different familiarity to the treadmill between the exercise and control group. The walking wheel (Model 80801) had an internal diameter of 15 cm and width of 5.7 cm, and was equipped with a safety mesh netting (Model 80801MSH25) to prevent the animal’s tail from being pinched. Animals were brought to the experimental suite 16 hours before mapping experiments and were fasted of food overnight with water *ad libitum*.

For 2DG uptake, the animal was administered IP [^14^C]-2DG (cat # MC355, Moravek Inc., Brea, CA, USA) at 0.3□µCi/g bodyweight in 0.53□ml normal saline. The animal was subsequently placed inside the closed walking wheel to walk at 2 m/min for 60 min to allow uptake of the tracer. At the end of walking, the animal was euthanized by cervical dislocation and the brain was extracted and flash-frozen in methylbutane over dry ice (−55□°C). The brains were later serially sectioned into 20-μm coronal slices, sampled with a 140-μm inter-slice distance, in a cryostat at -18□°C (Mikron HM550 OMP, Thermo Fisher Scientific, Waltham, MA, USA). Slices were heat-dried on glass slides and exposed to Kodak Biomax MR diagnostic film (Eastman Kodak, Rochester, NY, USA) for 3 days at room temperature. Autoradiographs were then digitized on an 8-bit grey scale using a voltage-stabilized light box (Northern Light R95 Precision Illuminator, Imaging Research Inc., St. Catharines, Ontario, Canada) and a Retiga 4000R charge-coupled device monochrome camera (QImaging of Teledyne Photometrics, Tucson, AZ, USA).

### 2.5. Whole-brain analysis of regional cerebral glucose uptake

For each animal, a 3D brain was reconstructed from 66 digitized, autoradiographic images of coronal sections (voxel size: 40 × 140 × 40 μm^3^) using our prior methods (Nguyen et al., 2004). Sections were selected starting at +2.4 mm anterior to the internal landmark of bregma. Adjacent sections were aligned using TurboReg, an automated pixel-based registration algorithm implemented in ImageJ (v.1.35, https://imagej.nih.gov/ij/index.html). This algorithm registered each section sequentially to the previous section using a non-warping geometric model that included rotations, rigid-body transformation and nearest-neighbor interpolation. We and others have adapted the SPM package (Wellcome Centre for Neuroimaging, University College London, London, UK) for the analysis of rodent autoradiographic cerebral blood flow and CGU data. For preprocessing, one mouse brain was selected as reference. All brains were spatially normalized to the reference brain in SPM (version 5). Spatial normalization consisted of applying a 12-parameter affine transformation followed by a nonlinear spatial normalization using 3D discrete cosine transforms. All normalized brains were then averaged to create a final brain template. Each original brain was then spatially normalized to the template. Final normalized brains were smoothed with a Gaussian kernel (full-width at half-maximum□=□ 240 × 420 × 240□μm^3^) to improve the signal-to-noise ratio. Proportional scaling was used to scale the voxel intensities so that the whole-brain average CGU was the same across animals. Global cerebral glucose uptake is believed to change very little with normal physiological alterations in cerebral functional activity (Sokoloff, 1991). This would be expected during the slow walking task in this study.

Unbiased, voxel-by-voxel Student’s *t-*tests between the exercise and control group were performed across the whole brain to access changes in rCGU following exercise using SPM. Threshold for statistical significance was set at *P*□<□0.05 at the voxel level with an extent threshold of 200 contiguous significant voxels. This combination reflected a balanced approach to control both Type I and Type II errors. The minimum cluster criterion was applied to avoid basing our results on significance at a single or a small number of suprathreshold voxels. Brain regions were identified according to mouse brain atlases (Dong, 2008; Franklin and Paxinos, 2008). Color-coded functional overlays showing statistically significant changes in rCGU were displayed over coronal sections of the template brain in MRIcro (v.1.40, https://people.cas.sc.edu/rorden/mricro/mricro.html).

### 2.6. Functional connectivity analysis of the cortico-basal ganglia-thalamic-cortical network

We took an ROI approach to assessing brain functional connectivity. A total of 176 ROIs were defined for six structures critical to motor and cognitive functions, including the CP, SNr, GPe, prefrontal cortex (PFC, including infralimbic, IL; prelimbic, PrL; cingulate area 1 and 2, Cg1 and Cg2), motor cortex (including primary and secondary motor, M1 and M2), and thalamic nuclei (anterodorsal, AD; anteromedial, AM; anteroventral, AV; central medial, CM; mediodorsal, MD; ventral anterior/ventrolateral, VA/VL; ventromedial, VM) (**Fig. 2**). ROIs were drawn on coronal sections of the template brain in MRIcro. A group of ROIs were defined for each structure at a given bregma level, e.g. 5 ROIs were defined for the rostral caudoputamen at bregma + 1.3 mm (CPr + 1.3, **Fig 2A**). ROIs for CP, SNr, and GPe were based on mesoscopic-level domain definitions as set forth in the mouse brain anatomical connectome (Hintiryan et al., 2016; Foster et al., 2021). Domain definition maps were transcribed to a visual template. Overlay of this template on to the digitized images allowed ROI definition in a standardized manner. Circular ROIs were drawn near the approximate center of each domain. Some domains too small in size in the SNr and GPe were not included in this analysis. ROIs for PFC, motor cortex, and thalamus were based on the mouse brain atlas (Dong, 2008; Franklin and Paxinos, 2008). A second set of brain slices collected adjacent to each autoradiographic section were histochemically stained for cytochrome oxidase. These histochemical images showing cytoarchitectural details were used to assist in the brain area identification in the autoradiographic images.

**Figure 1.**
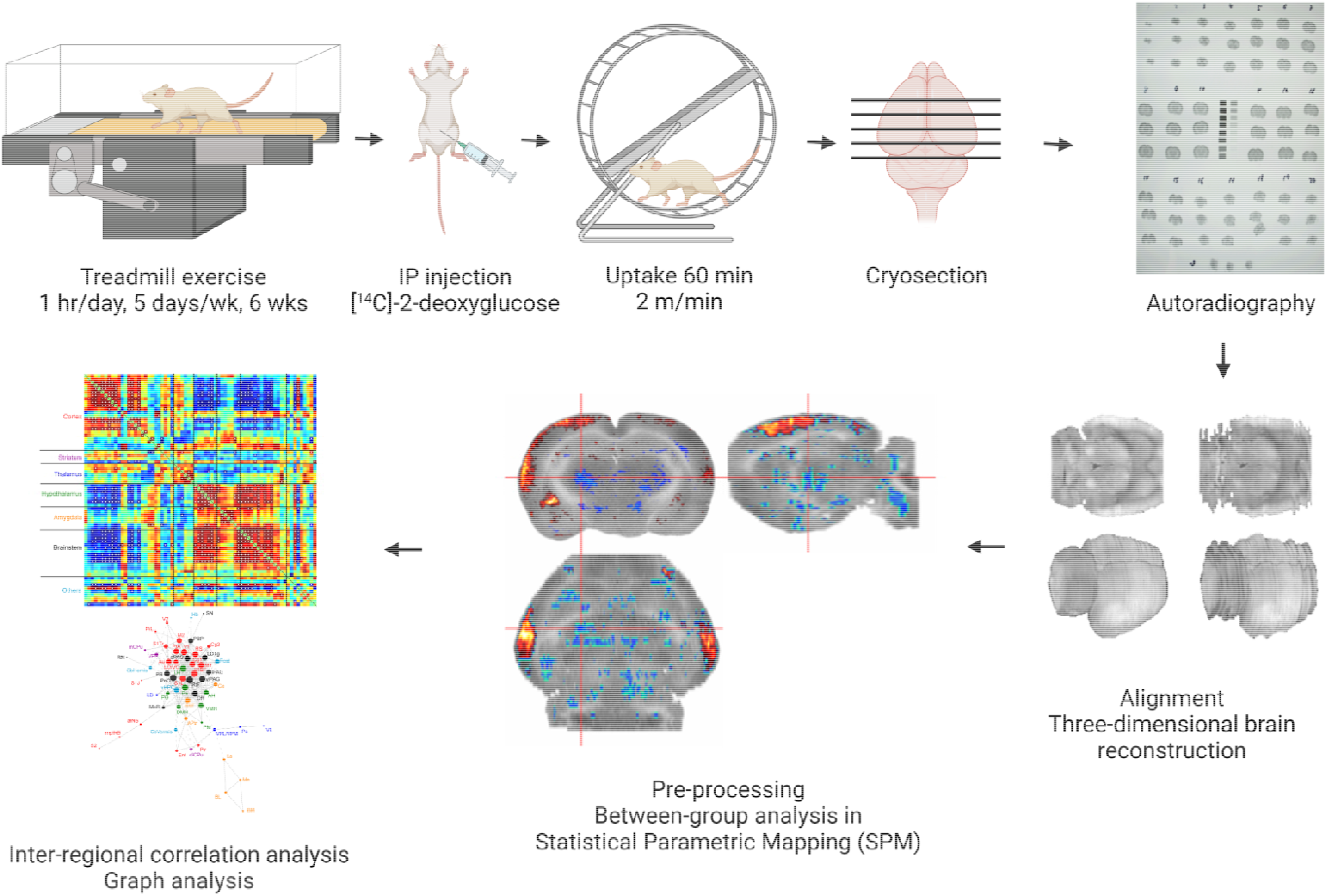
Experiment protocol. Created with BioRender.com.

**Figure 2.**
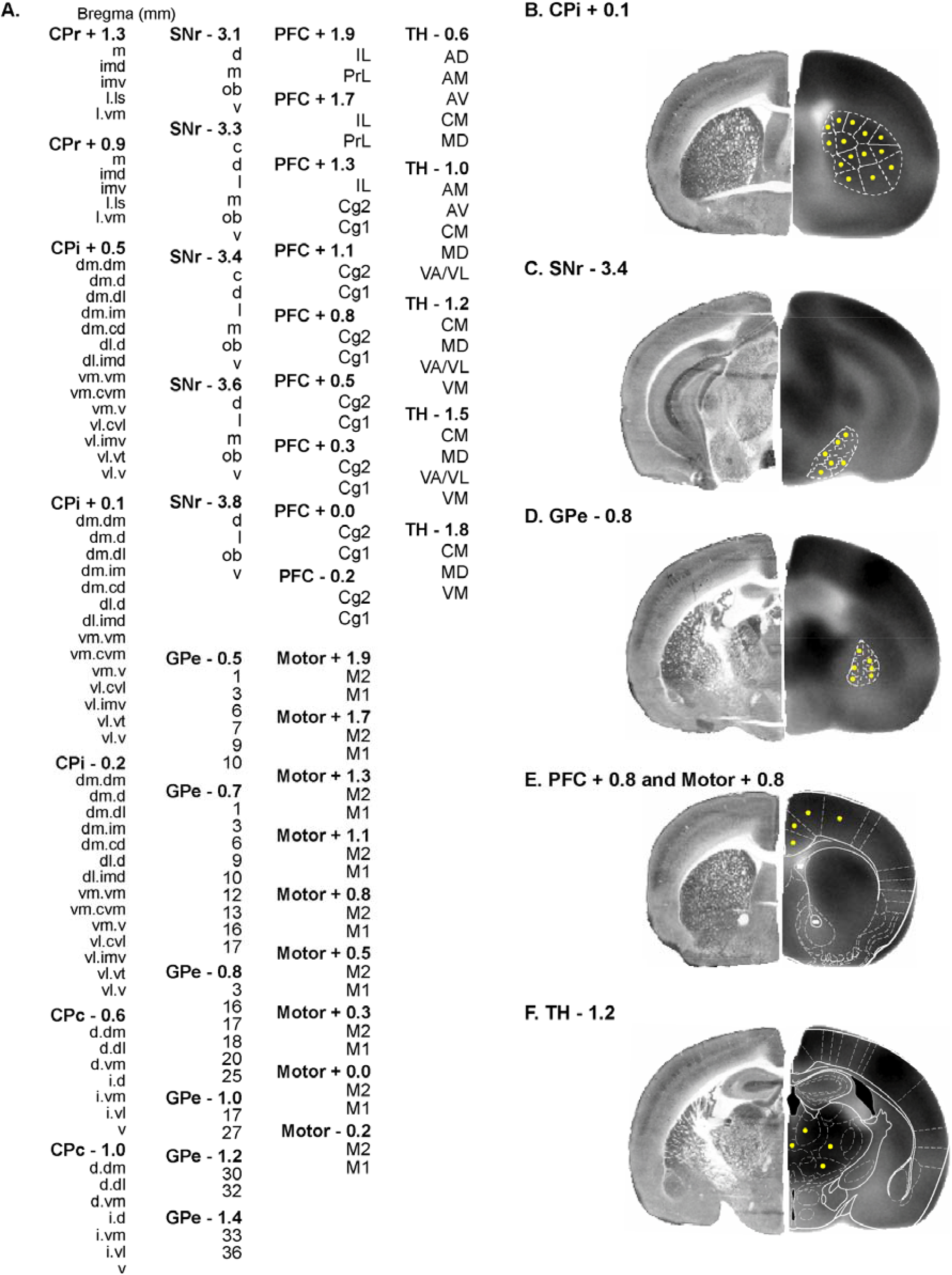
Definition of regions of interest (ROIs) for functional connectivity analysis. **(A)** List of 176 ROIs defined. A group of ROIs are defined for each structure at a given bregma level, e.g. CPr + 1.3, rostral caudoputamen at bregma + 1.3 mm. CPi/CPc, intermediate/caudal caudoputamen; SNr, substantia nigra pars reticulata; GPe, globus pallidus externus; PFC, prefrontal cortex (Cg1/Cg2, cingulate area 1/2. IL, infralimbic. PrL, prelimbic); M1/M2, primary/secondary motor cortex; TH, thalamic nuclei (AD, anterodorsal. AM, anteromedial. AV, anteroventral. CM, central medial. MD, mediodorsal. VA/VL, ventral anterior/ventrolateral. VM, ventromedial). **(B)** ROI definition for CPi domains at bregma + 0.1 mm as defined in (Hintiryan et al., 2016). The hemisphere on the left shows histochemical staining for cytochrome oxidase from a representative brain used to assist brain area identification. The hemisphere on the right is a coronal section of the template brain showing [^14^C]-2-deoxyglucose uptake. ROIs (red circles) are drawn near the approximate center of each domain. **(C)(D)** ROI definition for SNr domains at bregma -3.4 mm and GPe domains at -0.8 mm according to Foster et al. (2021). Some domains too small in size were not included in this analysis. **(E)** ROI definition for PFC and motor cortex at bregma + 0.8 mm. White outlines are modified from the mouse brain atlas (Franklin and Paxinos, 2008). **(F)** ROI definition for TH at bregma -1.2 mm.

Mean optical density of each ROI was extracted from each mouse brain using the MarsBaR toolbox for SPM (v.0.42, http://marsbar.sourceforge.net). We applied pairwise inter-regional correlation analysis to investigate brain functional connectivity. This is a well-established method, which has been applied to analyze rodent brain mapping data of multiple modalities, including autoradiographic 2DG (Soncrant et al., 1986; Nair and Gonzalez-Lima, 1999), autoradiographic cerebral blood flow (Wang et al., 2011), cytochrome oxidase histochemistry (Shumake et al., 2004; Fidalgo et al., 2011), activity regulated c-*fos* gene expression (Wheeler et al., 2013), and functional magnetic resonance imaging (fMRI) data (Schwarz et al., 2007). In this approach, correlations were calculated in an *inter*-subject manner at a single time point, i.e., across subjects within a group, and similar to functional connectivity analyses often performed in positron emission tomography (PET) data. The method precluded analysis of temporal dynamics of functional brain activation and differed from the *intra*-subject cross correlation analysis often used on fMRI time series data. While these different brain mapping modalities and analytic methods provide complementary information on brain functional connectivity, in comparing the results one should consider the possibility that differences in the time scales of data sampling, may result in the differential recruitment of ancillary regions (Di and Biswal, 2012; Buckner et al., 2013; Honey et al., 2007; Hutchison et al., 2013; Wehrl et al., 2013).

Pearson’s correlation coefficients between pairs of ROIs were calculated across subjects within a group in Matlab (Mathworks, Inc., Natick, MA, USA) to construct a correlation matrix. The matrices were visualized as heatmaps with Z-scores of Pearson’s correlation coefficients color-coded. Statistical significance of between-group difference in correlation coefficient was evaluated using the Fisher’s Z-transform test (*P* < 0.05)

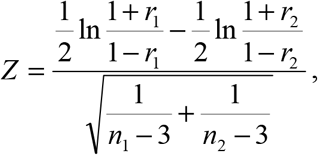

where *r*_1_ and *r*_2_ denote correlation coefficient in group 1 and group 2, while *n*_1_ and *n*_2_ denote sample size for each group. A positive Z value indicates that *r*_1_ is greater than *r*_2_.

To further facilitate between-group comparison of intra-or inter-structural functional connectivity, we defined *connectivity density* for a structure (or between two structures) as the number of connections expressed as a percentage of the total number of possible connections. For example, the total number of possible intra-structural connections among the 18 motor cortex ROIs is 153. There were 45 positive intra-motor cortex connections in the exercise group. The intra-motor cortex connectivity density was therefore + 29.4%. Connectivity density was calculated separately for positive and negative connections (correlations).

To delineate organization of the functional networks identified by the correlation matrices, graph theoretical analysis was performed as previously described (Wang et al., 2012) with the Pajek software (version 2.03, http://pajek.imfm.si/doku.php). Each ROI was represented by a node in a graph, and two nodes with significant correlation (positive or negative) were linked by an edge. A Kamada–Kawai algorithm was implemented to arrange (energize) the graph such that strongly connected nodes were placed closer to each other, while weakly connected nodes were placed further apart. Such energized graph provided an intuitive visualization of the network organization. To identify network hubs, connectivity degree of each node (degree centrality) was calculated as the number of edges linking it to the rest of the network. Intuitively, nodes with higher degrees were more central in the network organization. Nodes with degrees ranked in the top 10% were considered hubs.

### 2.7. Cytochrome oxidase histochemical staining

A second set of brain slices adjacent to sections of the autoradiographic reference brain were collected and histochemically stained for cytochrome oxidase. Histochemical images showing cytoarchitectural details were used to assist brain area identification in the autoradiographic images. Histochemical staining was undertaken with a protocol adapted from (Puga et al., 2007). In brief, staining proceeded at 4 °C as follows: (a) Pre-incubation fixation for 5 min in a phosphate buffer (0.1 M, pH 7.6) containing 10% sucrose and 0.5% glutaraldehyde; (b) Rinse 5 min x 3 times with phosphate buffer (0.1 M, pH 7.6) containing 10% sucrose; (c) Color intensification for 10 min in a Tris buffer (0.05M, pH 7.6) containing 275 mg/L cobalt chloride (CoCl_2_), 0.5% DMSO, and 10% sucrose; (d) Rinse for 5min with phosphate buffer (0.1 M, pH 7.6) containing 10% sucrose; (e) Staining incubation for 60 min at 37 °C with O_2_ bubbling in 700 ml 0.1M phosphate buffer containing 10% sucrose, 14 mg of catalase, 350 mg of diaminobenzedine tetrahydrochloride (DAB), 52.5 mg cytochrome c, and 1.75 ml of DMSO; (f) Stain termination/fixation at RT for 30 min in 0.1M phosphate buffer containing 10% sucrose and 10% formalin; (g) Dehydration with ethanol and clearing with xylene. Slides were coverslipped with Permount. Histological images were digitized and used to reconstruct a 3D brain as described above for autoradiographic images.

## 3. RESULTS

### 3.1. Effects of exercise on regional cerebral glucose uptake

Exercise resulted in broad changes across the CBT network during wheel walking (**Fig. 3**). Significant rCGU decreases were seen in the exercise compared to control group in motor regions, including primary motor cortex, the basal ganglia (intermediate CP, SNr), zona incerta, cerebellum (vermis, crus 2 of the ansiform lobule), as well as associated motor regions (cuneiform nucleus, precuneiform area), and sensory regions, including cortical areas (auditory, IL, primary somatosensory), medial geniculate nucleus, inferior colliculus, dorsal and ventral cochlear nucleus, anterior pretectal nucleus, and pontine reticular nucleus oral part. Exercised compared to control animals showed statistically significant rCGU increases in the limbic areas, including the hippocampus (CA1, CA2, CA3 fields, dentate gyrus, fimbria), parasubiculum, entorhinal cortex, piriform cortex, insula, amygdala, hypothalamus (lateral and ventromedial), nucleus accumbens, dorsal raphe, periaqueductal gray (ventrolateral, supraoculomotor) (*P* < 0.05, ≥ 200 significant contiguous voxels). Significant increases were also seen in the secondary somatosensory, primary and secondary visual, perirhinal, parietal association, and temporal association cortices, as well as in the olfactory tubercle, reticular thalamic nucleus, habenular nucleus, superior colliculus, and laterodorsal tegmental nucleus.

**Figure 3.**
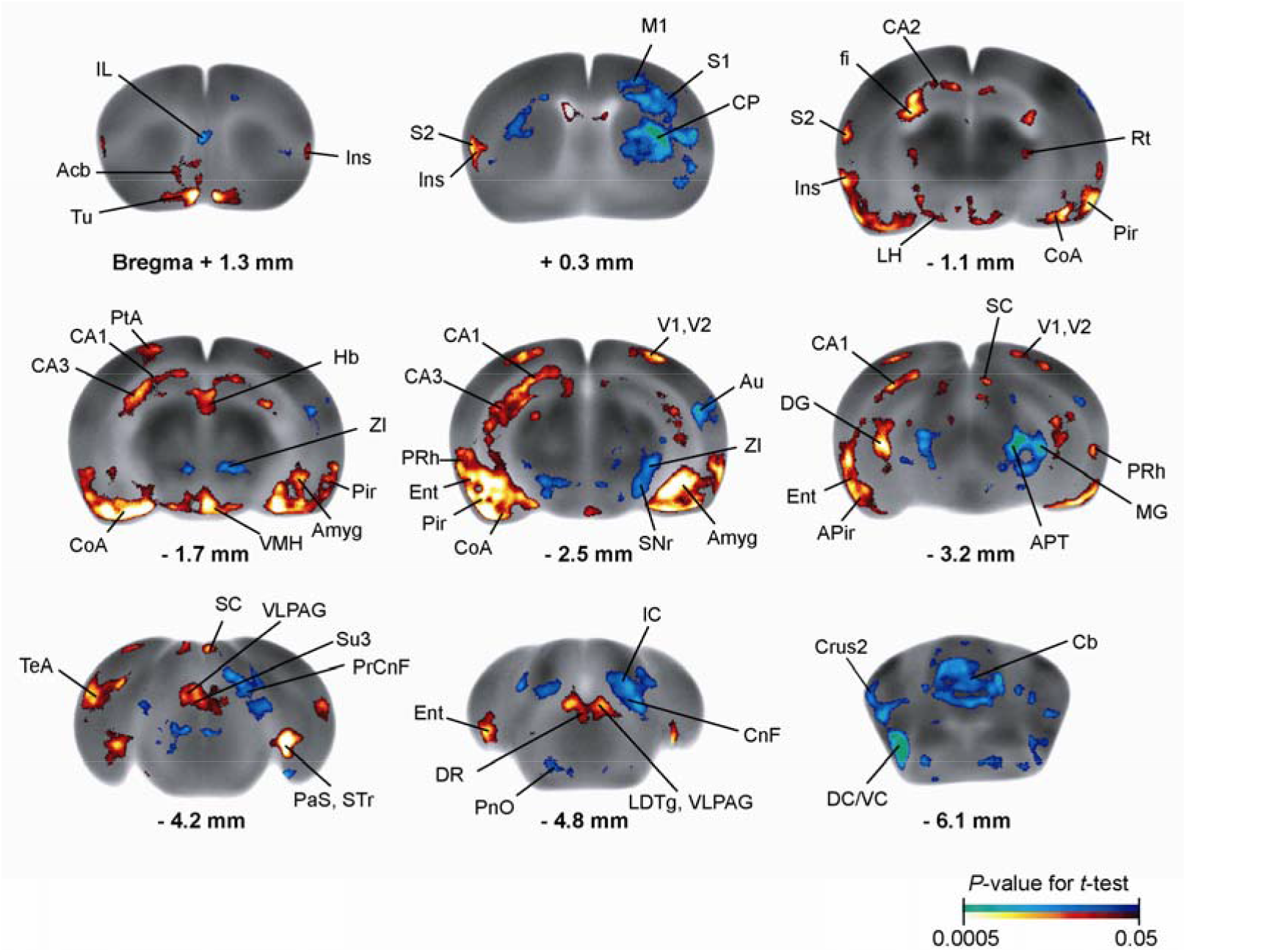
Exercise effects on regional cerebral glucose uptake (rCGU). Color-coded overlays show statistically significant increases (red) and decreases (blue) in rCGU in the exercise compared to the control group (*P* < 0.05 and extent threshold > 200 contiguous voxels, Student’s *t*-test). Shown are representative coronal sections of the template brain. Regions are identified according to the mouse brain atlas (Dong, 2008; Franklin and Paxinos, 2008). Abbreviations: Acb, nucleus (n.) accumbens; Amyg, amygdala; APir, amygdalopiriform transition; APT, anterior pretectal n.; Au, auditory cortex (cx); CA1/CA2/CA3, field CA1/CA2/CA3 hippocampus; CnF, cuneiform n.; CoA, cortical amygdala; CP, caudoputamen; Crus2, crus 2 of the ansiform lobule; DC/VC, dorsal/ventral cochlear n.; DG, dentate gyrus; DR, dorsal raphe n.; Ent, entorhinal cx; fi, fimbria; Hb, habenular n.; IC, inferior colliculus; IL, infralimbic cx; Ins, insular cx; LDTg, laterodorsal tegmental n.; LH, lateral hypothalamus; M1, primary motor cx; MG, medial geniculate n.; PaS, parasubiculum; Pir, piriform cx; PnO, pontine reticular n. oral part; PrCnF, precuneiform area; PRh, perirhinal cx; PtA, parietal association cx; Rt, reticular thalamic n.; S1/S2, primary/secondary somatosensory cx; SC, superior colliculus; SNr, substantia nigra pars reticulata; STr, subiculum transition area; Su3, supraoculomotor periaqueductal gray; TeA, temporal association cx; Tu, olfactory tubercle; V1/V2, primary/secondary visual cx; VLPAG, ventrolateral periaqueductal gray; VMH, ventromedial hypothalamic n.; ZI, zona incerta.

### 3.2. Sedentary control group: Functional connectivity of the cortico-basal ganglia-thalamic network

In the control group (**Fig. 4A and Supplementary Table S1**), functional connectivity of the network was characterized by strong, positive intra-structural connectivity in the SNr (+ 45.7%, positive connectivity density), GPe (+ 58.1%), and thalamus (+ 48.6%); and modest, primarily positive intra-structural connectivity in the CP (+ 11.0%), motor cortex (+ 23.5%), and mPFC (+ 19.3%) (**Fig. 4A**, along the diagonal line). Primarily positive inter-structural connectivity was seen between the CP and GPe (+ 13.9%), motor cortex and PFC (+ 9.6%), motor cortex and thalamus (+ 12.7%), and PFC and thalamus (+ 17.0%). The PFC showed primarily negative connectivity with the basal ganglia: with CP (−7.8%, negative connectivity density), with SNr (−7.4%), and with GPe (−21.4%) (**Supplementary Table S2**).

**Figure 4.**
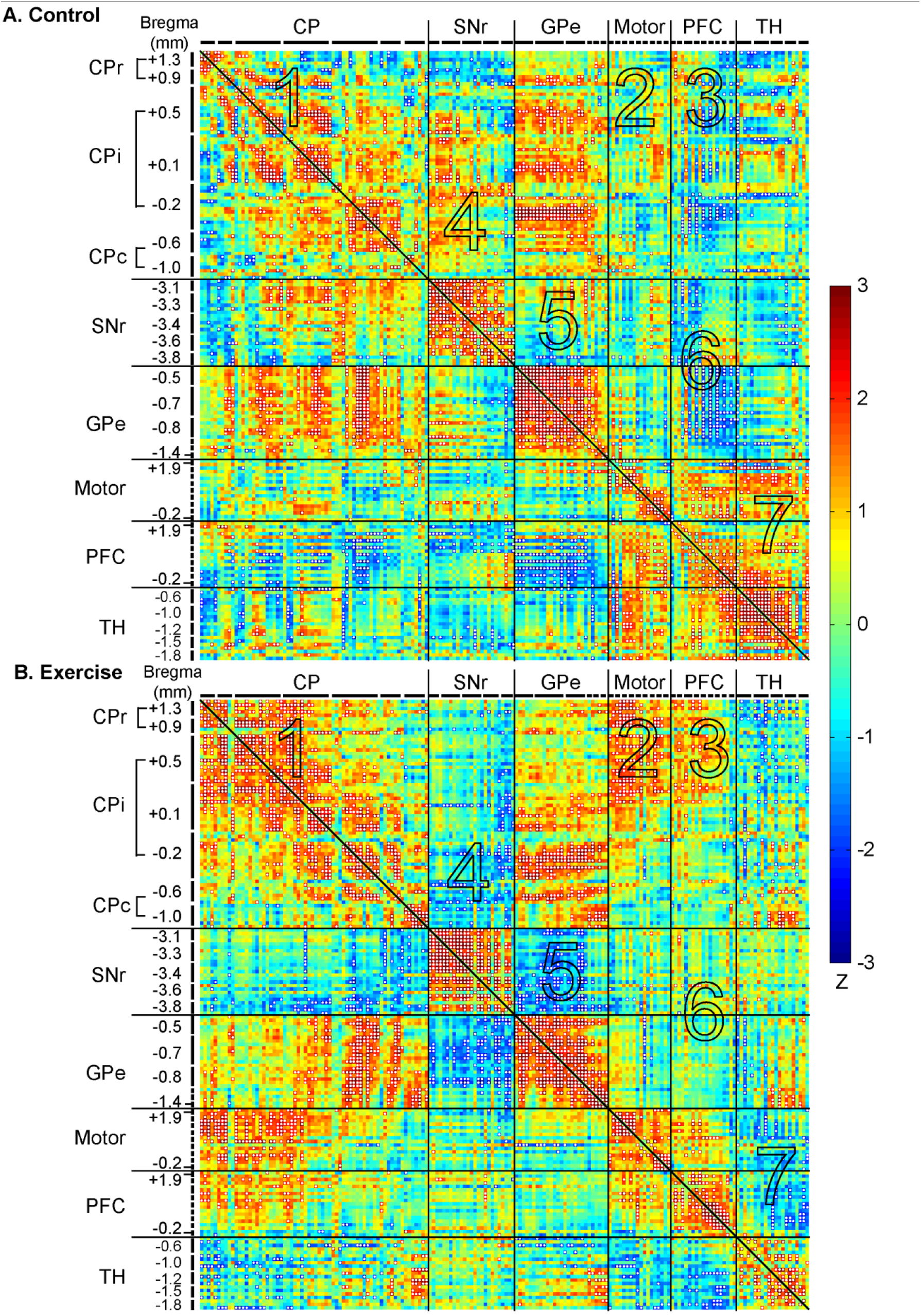
Exercise effects on functional connectivity of the cortico-basal ganglia-thalamic-cortical network. **(A)** Inter-regional correlation matrix shows functional connectivity patterns in the control group. Z scores of Pearson’s correlation coefficients are color-coded with positive and negative values shown in warm and cold colors, respectively. The matrix is symmetric across the diagonal line from upper left to lower right. Significant correlations (*P* < 0.05) are marked with white dots and interpreted as functional connections. Regions of interest are arranged in the sequence shown in **Fig. 2A**. They are grouped by structure, and further grouped by bregma level (marked by short, black lines) and arranged from rostral to caudal within each structure. **(B)** Inter-regional correlation matrix in the exercise group. CP, caudoputamen (CPr/CPi/CPc, rostral/intermediate/caudal); SNr, substantia nigra pars reticulata; GPe, globus pallidus externus; PFC, prefrontal cortex; TH, thalamus. The large numbers embedded in the heatmaps label pathways showing major exercise effects: 1, intra-CP; 2, CP-Motor cortex; 3, CP-PFC; 4, SNr-CP; 5, SNr-GPe; 6, PFC-SNr and PFC-GPe; 7, intra-TH, TH-Motor cortex, and TH-PFC.

**Fig. 5A** shows a connectivity graph of the control group based on the correlation matrix. The graph was energized using the Kamada-Kawai algorithm to help visualize network organization. Consistent with their high intra-structural connectivity, the SNr (blue nodes), GPe (green), and thalamus (white) each formed separate clusters. The motor cortex nodes (black) were closely connected with the thalamic cluster. The CP nodes (red) and PFC nodes (yellow) were both more scattered, making connections with all other clusters, while showing a particularly high level of integration with the GPe cluster. Nodes with the highest connectivity degrees (top 10%) were considered network hubs and included 12 GPe nodes, 3 intermediate CP (CPi) nodes, and 3 PFC nodes (Cg2).

**Figure 5.**
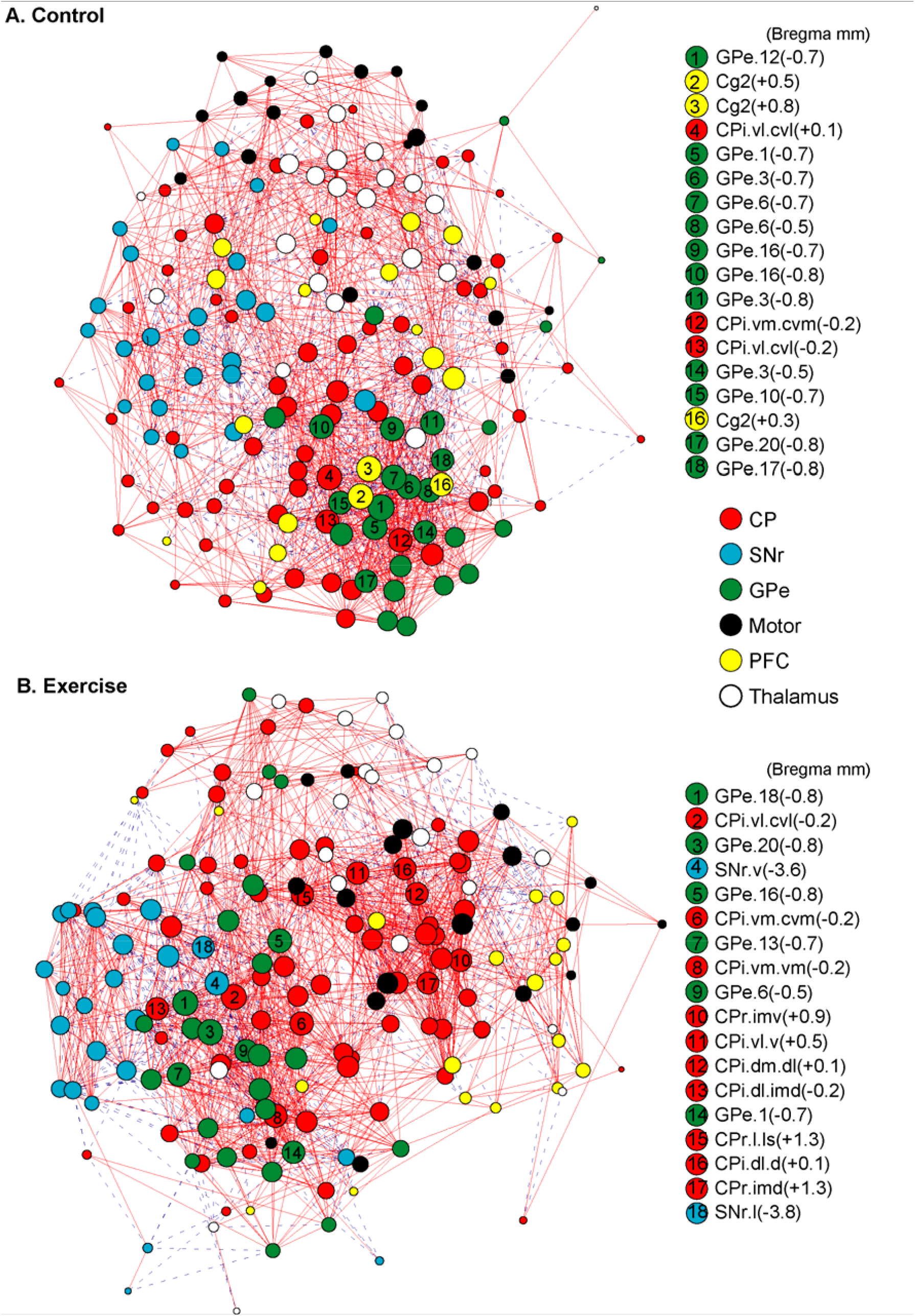
Exercise effects on the organization of connectivity graphs of the cortico-basal ganglia-thalamic-cortical network. **(A)** The functional connectivity network of the control group is represented with a graph, in which nodes represent regions of interest (ROIs) and edges represent significant correlations. Solid red lines denote significant positive correlations, whereas dashed blue lines denote significant negative correlations. The graph is energized using the Kamada–Kawai algorithm that places strongly connected nodes closer to each other while keeping weakly connected nodes further apart. The size of each node (in area) is proportional to its degree, a measurement of the number of connections linking the node to other nodes in the network. ROIs with the highest degree (top 10%) are considered hubs of the network and labeled with their ranking numbers. Nodes are color-coded to facilitate identification of nodes belonging to the same structure.(**B**)Connectivity graph of the exercise group. CP, caudoputamen; SNr, substantia nigra pars reticulata; GPe, globus pallidus externus; PFC, prefrontal cortex.

### 3.3. Exercise group: Functional connectivity of the cortico-basal ganglia-thalamic network

The exercise compared to control group showed broad changes in functional connectivity (**Fig. 4B, Fig. 6, and Supplementary Tables S1 and S2**) and network organization (**Fig. 5B**). The following major differences comparing exercise and control group correspond to areas numbered 1 through 7 in **Fig. 4** and **Fig. 6**. (1) Intra-CP connectivity density increased from + 11.0% in the control to + 20.0% in the exercise group. Increased connections were mainly located in rostral CP (CPr) and dorsal aspect of CPi. (2) In the exercise group, new, positive connections were formed between CPr/CPi and motor cortex. (3) CP-PFC connectivity changed polarity from -7.8% in the control to + 3.6% in the exercise group. New, positive connections involved mainly CPr and CPi. (4) CP-SNr connectivity turned predominantly negative in the exercise group, with a -6.3% density involving mainly CPi and caudal CP (CPc). (5) SNr-GPe connectivity turned predominantly negative in the exercise group, with a -18.5% density. (6) Negative connectivity between PFC and the basal ganglia observed predominantly in the control group was largely absent in the exercise group. (7) Intra-thalamic connectivity density decreased from + 48.6% in the control to + 22.4% in the exercise group. A predominantly positive thalamus-motor cortex connectivity (+ 12.7%) and thalamus-PFC connectivity (+ 17.0%) seen in the control group changed polarity in the exercise group to a predominantly negative connectivity of -5.3% and -7.3%, respectively.

**Figure 6.**
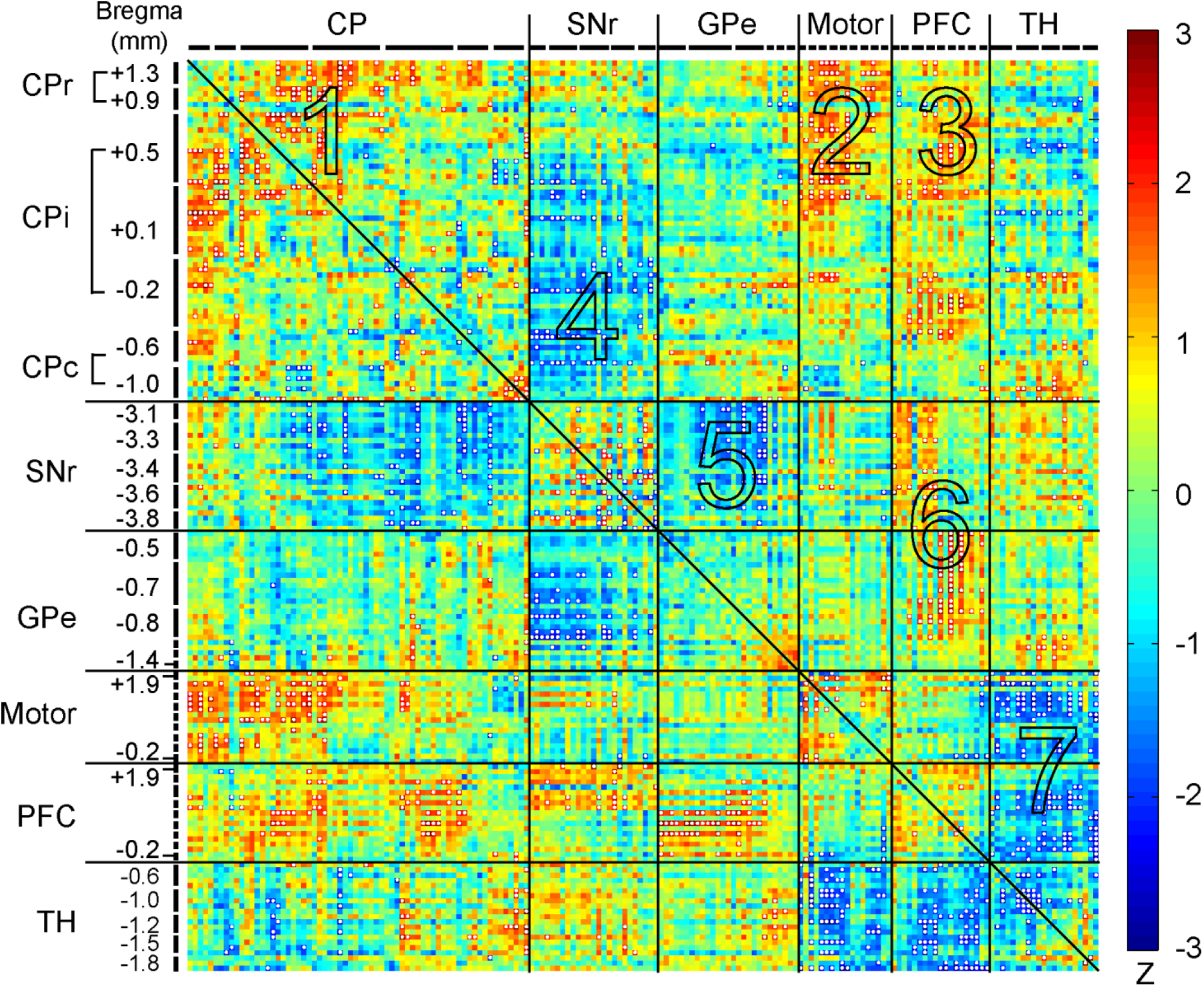
Exercise vs. control group: Changes in functional connectivity. The matrix of Fisher’s Z-statistics represents differences in Pearson’s correlation coefficients (*r*) between exercise and control groups. Positive/negative Z values indicate greater/smaller *r* in the exercise compared to control group. Significant between-group differences (*P* < 0.05) were marked with white dots. CP, caudoputamen (CPr/CPi/CPc, rostral/intermediate/caudal); SNr, substantia nigra pars reticulata; GPe, globus pallidus externus; PFC, prefrontal cortex; TH, thalamus. The large numbers embedded in the heatmap label pathways showing major exercise effects: 1, intra-CP; 2, CP-Motor cortex; 3, CP-PFC; 4, SNr-CP; 5, SNr-GPe; 6, PFC-SNr and PFC-GPe; 7, intra-TH, TH-Motor cortex, and TH-PFC.

**Fig. 5B** shows energized connectivity graph of the exercise group. The PFC nodes (yellow) and thalamic nodes (white) were marginalized. Motor cortex nodes (black) became more integrated towards the center of the network through connections with CP nodes (red). The CP played a more central role in the network, with a greater number of CP nodes functioning as hubs compared to in the control group. These CP nodes showed greater connectivity with the GPe and SNr through ROIs mostly caudal to the bregma, and with the motor cortex through ROIs mostly rostral to the bregma. Network hubs consisted of 10 CP nodes (up from 3 in the control group), 6 GPe nodes (down from 12 in the control group), and 2 new SNr nodes. There was no network hub in the PFC in the exercise group, consistent with decreases in functional connectivity of the PFC nodes with the other structures (**Supplemental Table S2**).

**Fig. 7** shows connectivity degree changes in the exercise compared to control group of all ROIs in a ranked order. Among those with the highest gains in degree were CPr, CPi, SNr, M1, and M2 ROIs (**Fig. 7A**). ROIs with the greatest losses in degree were from the Cg2, thalamus, and GPe (**Fig. 7B**).

**Figure 7.**
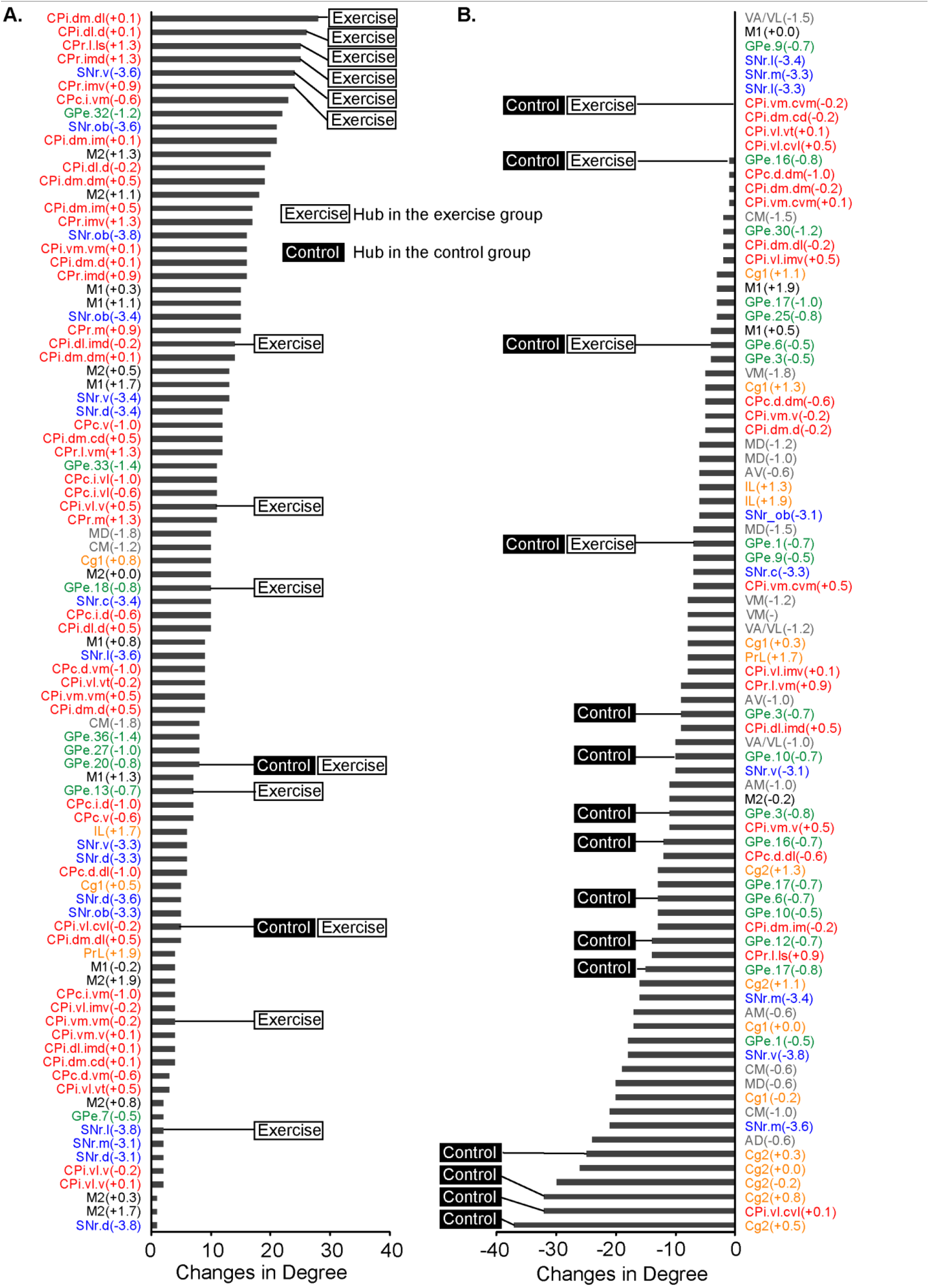
Functional connectivity degree changes comparing the exercise and control group. **(A)** Regions of interest (ROIs) showing increases in degree in the exercise compared to control group. **(B)** ROIs showing no changes or decreases in degree in the exercise compared to control group. Bregma levels of ROIs are included in parentheses. ROIs identified as network hubs in **Figs. 5A and 5B** are labeled with “Control” and “Exercise” text boxes. CPr/CPi/CPc, rostral /intermediate/caudal caudoputamen; SNr, substantia nigra pars reticulata; GPe, globus pallidus externus; Cg1/Cg2, cingulate cortex area 1/2; IL, infralimbic cortex; PrL, prelimbic cortex; M1/M2, primary/secondary motor cortex; thalamic nuclei (AD, anterodorsal. AM, anteromedial. AV, anteroventral. CM, central medial.MD, mediodorsal. VA/VL, ventral anterior/ventrolateral. VM, ventromedial).

**Fig. 8** summarizes functional connectivity degrees of CP domains in the control and exercise groups, as well as exercise-associated changes along the rostral-caudal axis. The domain maps were modified from (Hintiryan et al., 2016). In the exercise compared to control group, domains showing the most gains in degree included most domains in the CPr, the dorsomedial area of CPi (CPi.dm), the dorsolateral area of CPi (CPi.dl.d), and the intermediate and ventral areas of CPc (CPc.i.vm, CPc.v), while some domains in the lateral area of CPr (CPr.l), ventral areas of CPi (CPi.vm, CPi.vl), and dorsal area of CPc (CPc.d) showed decreases in degree.

**Figure 8.**
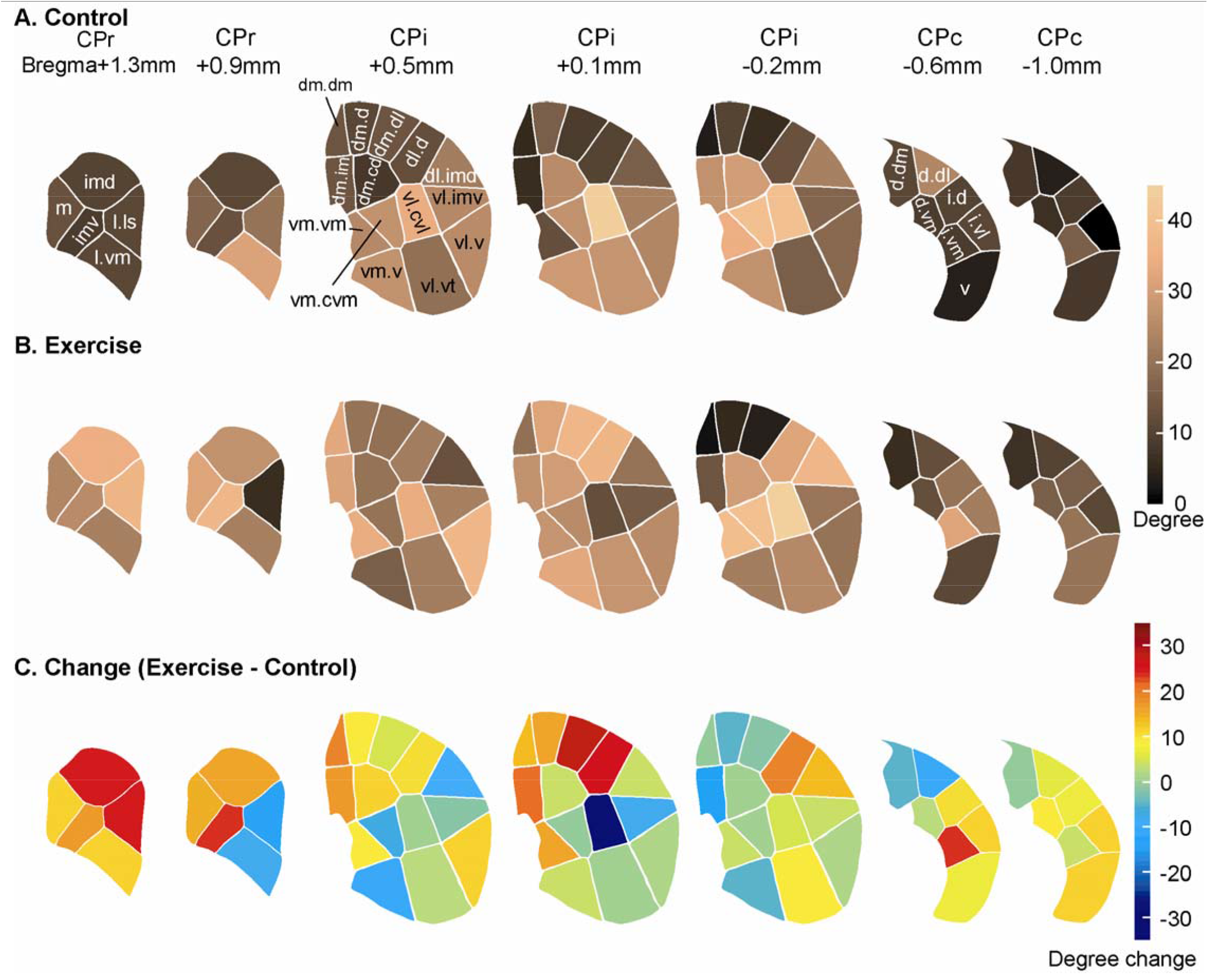
Functional connectivity degree changes in the caudoputamen (CP). **(A)** Connectivity degree of CP domains in the control group is color-coded. **(B)** Connectivity degree of CP domains in the exercise group. **(C)** Connectivity degree changes in the exercise compared to control group. CPr/CPi/CPc, rostral/intermediate/caudal caudoputamen. Domain maps and nomenclature were drawn based on Hintiryan et al. (2016).

**Fig. 9** summarizes functional connectivity degrees and exercise-associated changes in the GPe and SNr domains. The domain maps were modified from (Foster et al., 2021). Exercise induced overall decreases in degree in the GPe domains, except in the caudal area (GPe.32, GPe.33, **Fig. 9A-C**). In the SNr (**Fig. 9D-F**), exercise induced degree decrease rostrally in the oro-brachial, ventral, and central domains (SNr.ob, SNr.v, SNr.c) and caudally in the medial and ventral domains (SNr.m, SNr.v); while inducing degree increase in domains at intermediate and caudal levels (SNr.ob at -3.4 to -3.8mm, SNr.v at -3.4 and -3.6mm, SNr.d and SNr.c at -3.4mm).

**Figure 9.**
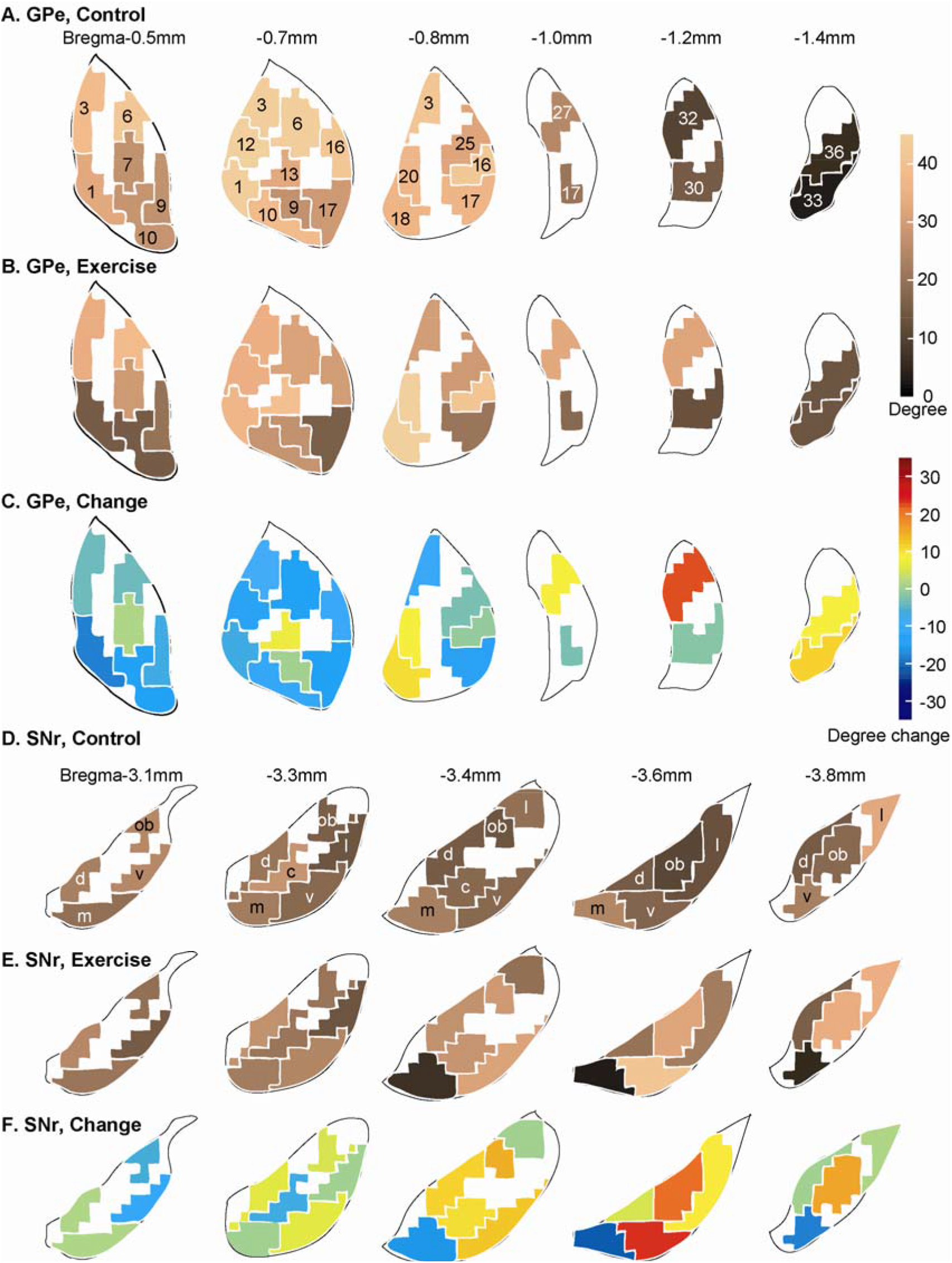
Functional connectivity degree changes in the globus pallidus externus (GPe) and substantial nigra pars reticulata (SNr). **(A)** Connectivity degree of GPe domains in the control group is color-coded. **(B)** Connectivity degree of GPe domains in the exercise group. **(C)** GPe connectivity degree changes comparing the exercise and control groups. **(D)** Connectivity degree of SNr domains in the control group. **(E)** Connectivity degree of SNr domains in the exercise group. **(F)** SNr connectivity degree changes comparing the exercise and control groups. Domain maps and domain nomenclature were drawn based on Foster et al. (2021).

## 4. DISCUSSION

We applied the 2DG autoradiographic cerebral metabolic mapping method to investigate exercise-associated functional reorganization in the normal mouse brain. Exercise significantly altered both regional cerebral glucose uptake in broad areas of the brain, as well as inter-regional functional interactions in the CBT network. Compared to the sedentary controls, the exercise group showed increases in positive functional connectivity within and between the CP and motor cortex, newly emerged negative connectivity of the SNr with GPe and CP, as well as diminished negative connectivity of the PFC with the CP. To our best knowledge, this is the first study that systematically analyzed functional connectivity in the CBT network at the mesoscopic level during learning of a new motor task. ROIs were chosen to conform to the subregional parcellation of the CP, SNr, and GPe in the mouse brain structural connectome. Using the structural connectome as a roadmap, the current study started to address network activity underlying the brain’s changes in learning capacity of a novel wheel walking task following chronic exercise.

### 4.1. Exercise-effects on regional cerebral glucose uptake

In our study, mice underwent chronic (six weeks), high-intensity exercise on a motorized horizontal treadmill or no exercise. Cerebral metabolic mapping was thereafter undertaken in all animals during a novel wheel walking challenge, with differences in rCGU of the CBT likely reflecting long-lasting cerebral functional reorganization. Exercise resulted in broad changes in rCGU during wheel walking, including decreases in the motor areas (primary motor cortex, dorsolateral aspect of intermediate CP, SNr, zona incerta, and the cerebellar vermis); but increases broadly in the limbic areas (the hippocampus, entorhinal cortex, amygdala, hypothalamus, piriform and insular cortex, dorsal raphe, periaqueductal gray and nucleus accumbens), as well as the visual and association cortices (parietal, temporal), and dorsolateral tegmental nucleus. This general pattern of changes was remarkably similar to what we previously observed in a cerebral blood flow (CBF) mapping study (Holschneider et al., 2007). In this earlier study, rats received 6 weeks of rotarod exercise and were subsequently imaged during a locomotor challenge. Compared to sedentary controls, animals exercised on the rotarod showed decreases in regional CBF in the motor pathway (M1, M2, dorsolateral CP, zona incerta, cerebellar vermis) and primary somatosensory cortex, as well as increases in limbic regions (hippocampus, entorhinal cortex, periaqueductal gray, amygdala). Of note, the above-mentioned patterns observed after chronic exercise were generally opposite to those elicited during acute locomotion, where prior work has shown increases in rCGU (Vissing et al., 1996) and CBF (Nguyen et al., 2004) in motor regions (motor cortex, striatum, substantia nigra, cerebellum) and decreases in rCGU in limbic regions (including amygdala, hippocampus, hypothalamus, dorsal raphe). This converging evidence across species and across brain mapping modalities supports a general pattern of exercise effects on functional cerebral reorganization.

Similar effects on the motor regions have been reported after motor training in humans when comparing professional musicians and novices during the performance of finger sequences (Munte et al., 2002). The magnitude of fMRI BOLD signals to simple, overpracticed finger tasks in experts was attenuated relative to that seen in novices in the motor cortex, basal ganglia, and cerebellar vermis (Jancke et al., 2000; Kim et al., 2004; Koeneke et al., 2004). In non-musicians, attenuation of activation in somatosensory and motor cortices has been reported as subjects become more practiced on finger tasks (Morgen et al., 2004). When finger sequences are pre-learned, a lesser and more circumscribed activation has been noted in the cerebellum (vermis and hemispheres) (Friston et al., 1992; Jenkins et al., 1994), and striatum (Tracy et al., 2001). These and our findings suggest that extensive motor training results in a functionally more efficient way to control movements. The nature of this increase in functional efficacy needs further investigation, but may involve a shift from anaerobic to aerobic metabolism (McCloskey et al., 2001; Garifoli et al., 2003; Navarro et al., 2004).

### 4.2. Exercise enhances functional connectivity of the caudoputamen and motor cortex

The notion that exercise enhanced the efficiency of the motor circuit received further support from the functional connectivity analysis. There was broadly increased intra- and inter-structural connectivity across the CP and motor cortex in exercised compared to control animals. The findings of increased functional connectivity in the CP and motor cortex in the exercised animals in the context of decreases or no change in rCGU in these regions suggests that the motor regions, rather than being deactivated, were functioning with greater integration at the network level. Such dissociation between regional activity and functional connectivity has been previously reported. Eisenstein et al. (2021) reported that in older adults a physically active lifestyle was associated with lower activity level of the hippocampus, but higher functional connectivity of the hippocampus to hubs of the default mode network during memory encoding. Conversely, Trujillo et al. (2015) reported that Parkinson’s patients compared to healthy controls showed increased activity but decreased functional connectivity in the dorsolateral prefrontal cortex during a visuospatial task. These and other findings (Pinho et al., 2014) together suggest that a simultaneous task-related decrease in regional activity and increase in functional connectivity may be a marker for high functionality of the region, while an increase in activity coupled with a decrease in connectivity may represent a marker for dysfunction.

In the exercise compared to the control group, the rostral level and dorsal aspect of intermediate level CP showed the greatest increase in functional connectivity. The CPr exhibits high integration among cortical afferents, across different cortical subnetworks, suggesting cross modality integration, while CPi.dm receives input from the medial cortical subnetwork (including visual, auditory, anterior cingulate, retrosplenial, and posterior parietal association areas) (Hintiryan et al., 2016). Increased functional connectivity in these CP areas suggests a shift in the CP subregional recruitment that facilitates cross modality integration.

### 4.3. Exercise decreases functional connectivity between prefrontal cortex and the basal ganglia

The prefrontal cortices project to the medial aspect of the CP and are implicated in cognitive functions, including executive function (O’Neill and Brown, 2007; Baker and Ragozzino, 2014; Grospe et al., 2018). While the PFC in the control group was functionally closely integrated with the basal ganglia through negative connectivity and with the motor cortex and thalamus through positive connectivity, it was functionally dissociated from these structures in the exercise group. The connectivity density of PFC with all other structures dropped from 12.6% in controls to 3.7% in the exercise group. It is believed that the PFC is critically involved in the early phase of a motor learning task (Dayan and Cohen, 2011). In fact, we have previously shown greater functional connectivity between the PFC and dorsomedial CP in rats during the early phase of learning a complex wheel walking task (Guo et al., 2017). As learning progresses, the PFC becomes less involved (Dayan and Cohen, 2011). Wheel walking in the current study was a new motor task for the animals. Both groups were habituated to the wheel for two days prior to the 2DG mapping experiments. The finding of high connectivity to outside structures (inter-structural connectivity) in the control group is consistent with the notion that in the early phase of motor skill learning, the PFC is critically involved in cognitive control. Reduced PFC connectivity to outside structures in the exercise group, coupled with an increased functional connectivity of the CP and motor cortex, suggests higher network efficiency that expedites motor skill learning and the transition during initial learning from PFC to motor cortex in terms of cortical recruitment.

### 4.4. Exercise led to strong negative functional connectivity between the SNr and GPe

In the basal ganglia, the CP and GPe were positively connected in both the control and exercise groups. In contrast, exercise induced substantial changes in the functional connectivity of SNr with CP and with GPe. In the control group, SNr-CP and SNr-GPe connectivity were relatively weak and contained both positive and negative connections. In the exercise group, SNr-CP and SNr-GPe turned exclusively negative to -6.3% and -18.5%, respectively. Negative connection (anticorrelation) is believed to reflect inter-regional modulation, possibly involving the suppression of excitability of a network (Gopinath et al., 2015). Though the direction of change related to excitatory-inhibitory inputs remains unresolved, negative correlations have been associated with known inhibitory connections in the rodent (Liang et al., 2012). In the indirect pathway of the basal ganglia, GPe inhibits SNr activity through inhibition of the subthalamic nucleus. The strong negative SNr-GPe connectivity, and decrease in rCGU in part of the SNr are consistent with greater activation of the indirect pathway in the exercise group.

### 4.5. Translational implications

An important clinical implication of our findings is that chronic exercise may prime the brain for accelerated new motor learning, whereby CP to motor cortex networks are heavily recruited compared to CP to PFC networks. Whereas traditional motor learning theories emphasize learning specific to the context and task performed through engagement of the PFC, recent studies suggest that a general, transferable knowledge about skill learning processes that involves the CP and motor cortex, may be acquired through prior motor learning (Seidler, 2004). The extent of generalization may depend on the breadth and duration of experience obtained, the degree of arousal (Loras et al., 2020), context, and intensity (Holman and Staines, 2021). Lehmann et al. (2020) showed that subjects who underwent cardiovascular exercise subsequently learned a dynamic balancing task faster compared to controls undergoing stretching. Exercise also induced increases in cerebral blood flow in frontal brain regions and changes in white matter microstructure in frontotemporal fiber tracts, suggesting a transfer potential of experience-induced brain plasticity. Inoue et al. (2018) showed that long-term exercise increased BDNF expression in the motor cortex and facilitated a transfer of motor learning from aerobic exercise to postural coordination. Aerobic exercise in stroke survivors improved cognitive domains related to motor learning (Quaney et al., 2009). It has been proposed that exercise-mediated improvements in motor learning can be mediated by discrete, experience-driven changes within specific neural representations subserving the performance of the trained task (Karni et al., 1998). However, few studies have examined the underlying functional reorganization of neural circuits. Our study highlights exercise-associated functional reorganization of the CBT circuit in areas implicated in cognitive and motor processing, which may mediate improved motor learning. Such circuit-level understanding may inform therapeutic use of exercise for the rehabilitation of patients with motor and cognitive dysfunctions.

### 4.6. The importance of systematic functional connectivity analysis at the mesoscopic level

Tremendous progress has been made in understanding the structural connectome of the rodent brain (Oh et al., 2014; Zingg et al., 2014; Bota et al., 2015; Hunnicutt et al., 2016; Knox et al., 2019). Understanding of the brain functional connectome remains much less advanced (Frégnac, 2017; Venkadesh and Van Horn, 2021). A large part of the challenge is that brain functional connectivity is dynamic and depends on not only the current behavioral state (task, context), but also past experience (learning and memory). Advances have been made to delineate the whole-brain level functional connectome using resting-state fMRI (rs-fMRI) (Stafford et al., 2014; Mills et al., 2018; Zerbi et al., 2019; Coletta et al., 2020; Yang et al., 2021) and at the system level for a specific behavior, such as conditioned fear recall (Wheeler et al., 2013; Holschneider et al., 2014). Fregnac (2017) has emphasized that understanding mesoscale (mesoscopic) organization and full network dynamics may reveal a simpler formalism than the microscale level. Our study selected basal ganglia ROIs based on novel, mesoscopic domain definitions in the mouse brain connectome. In addition, multiple ROIs were defined at different bregma levels for some CP, GPe, and SNr domains. We felt this sampling method, informed by state-of-the-art structural connectomic data, reflects the best effort (an optimal compromise) to delineate *functional units* within brain structures. Specifically, a subregional sampling may be needed to avoid losing information when signals are spatially averaged over, for instance, whole CP, globus pallidus, or motor cortex, as many prior 2DG studies have done. At the same time mesoscopic sampling provides sufficient data simplification, while avoiding the risk of losing relevance to the interpretation of behavior through an exhaustive reductionist analysis (Frégnac, 2017). In some cases, differences in functional connectivity were noted in the same domain across different bregma levels, e.g., CPr.l.vm (**Fig. 8**), suggesting the existence of multiple functional units within a domain. This may in turn inform further analysis of these domains in terms of gene expression, neurochemistry, and local circuitry.

### 4.7. Limitations

Our study focused on the CBT network, with an emphasis on the basal ganglia. Brain regions outside of the sampled CBT network may also contribute to the learning and performance of the wheel walking task and undergo changes in response to exercise. These regions include the hippocampus, cerebellum, parietal association cortex, somatosensory and visual cortices, and ventral striatum (nucleus accumbens, ventral pallidum), as shown in the activation map (**Fig. 3**). This limitation is due to the scope and aim of the study. Mesoscopic ROI selection for the hippocampus, cerebellum, and the cortical structures remains challenging due to the large size and incomplete understanding of subregional heterogeneity of these structures, and needs to be addressed in future work.

It is important to note that correlation is not causation. Interpretation of functional connectivity between two nodes, even with direct structural connectivity, is not trivial due to the existence of indirect pathways through other node(s), possible influence from a common third node, and reciprocal connections and loops common in neural networks. Nevertheless, this mapping method provides insight into exercise-induced functional network reorganization and informs further mechanistic and causal research that focus on specific brain regions or pathways by examining neuroplasticity and manipulation.

As noted above, the functional connectome represents a dynamic map. The time scales of data sampling matter. Exploring network structure of cerebral cortex on multiple time scales, Honey et al. (2007) reported that at slower time scales (minutes), the aggregate strength of functional connectivity between regions is, on average, a good indicator of the presence of an underlying structural link. At faster time scales, significant fluctuations are observed. Thus, while the aim of anatomic parcellation remains the discovery of discrete functional units, the answers provided may depend in part on the temporal resolution of data sampling.

### 4.8. Conclusion

Overall findings from our study support that exercise induced a significant functional reorganization of the CBT neural network that led to greater connectivity between the CP and motor cortex that may underlie gains in learning of a new motor task. Such findings support that exercise may facilitate motor learning through engagement of key motor networks important for the generalizability of motor performance and may be used to guide future rehabilitation programs.

## Acknowledgments

This work was supported by grants from the US Department of Defense (Army, CDMRP) grant # W81XWH18-1-0666 (DPH) and grant # W81XWH19-1-0443 (MWJ).

## REFERENCES

Baker PM, Ragozzino ME (2014) Contralateral disconnection of the rat prelimbic cortex and dorsomedial striatum impairs cue-guided behavioral switching. Learn Mem 21:368–379. DOI:10.1101/lm.034819.114.

Bota M, Sporns O, Swanson LW (2015) Architecture of the cerebral cortical association connectome underlying cognition. Proc Natl Acad Sci U S A 112:E2093–2101. DOI:10.1073/pnas.1504394112.

Buckner RL, Krienen FM, Yeo BT (2013) Opportunities and limitations of intrinsic functional connectivity MRI. Nat Neurosci 16:832–837. DOI:10.1038/nn.3423.

Coletta L, Pagani M, Whitesell JD, Harris JA, Bernhardt B, Gozzi A (2020) Network structure of the mouse brain connectome with voxel resolution. Sci Adv 6. DOI:10.1126/sciadv.abb7187.

Cotman CW, Berchtold NC (2002) Exercise: a behavioral intervention to enhance brain health and plasticity. Trends Neurosci 25:295–301. DOI:10.1016/s0166-2236(02)02143-4.

Dauwan M, Begemann MJH, Slot MIE, Lee EHM, Scheltens P, Sommer IEC (2021) Physical exercise improves quality of life, depressive symptoms, and cognition across chronic brain disorders: a transdiagnostic systematic review and meta-analysis of randomized controlled trials. J Neurol 268:1222–1246. DOI:10.1007/s00415-019-09493-9.

Dayan E, Cohen LG (2011) Neuroplasticity subserving motor skill learning. Neuron 72:443–454. DOI:10.1016/j.neuron.2011.10.008.

Di X, Biswal BB (2012) Metabolic brain covariant networks as revealed by FDG-PET with reference to resting-state fMRI networks. Brain Connect 2:275–283. DOI:10.1089/brain.2012.0086.

Dong HW (2008) The Allen reference atlas: A digital color brain atlas of the C57BL/6J male mouse. Hoboken, NJ, USA: John Wiley & Sons.

Duchesne C, Gheysen F, Bore A, Albouy G, Nadeau A, Robillard ME, Bobeuf F, Lafontaine AL, Lungu O, Bherer L, Doyon J (2016) Influence of aerobic exercise training on the neural correlates of motor learning in Parkinson’s disease individuals. Neuroimage Clin 12:559–569. DOI:10.1016/j.nicl.2016.09.011.

Eisenstein T, Giladi N, Hendler T, Havakuk O, Lerner Y (2021) Physically Active Lifestyle Is Associated With Attenuation of Hippocampal Dysfunction in Cognitively Intact Older Adults. Front Aging Neurosci 13:720990. DOI:10.3389/fnagi.2021.720990.

Fidalgo C, Conejo NM, Gonzalez-Pardo H, Arias JL (2011) Cortico-limbic-striatal contribution after response and reversal learning: a metabolic mapping study. Brain Res 1368:143–150. DOI:10.1016/j.brainres.2010.10.066.

Foster NN, Barry J, Korobkova L, Garcia L, Gao L, Becerra M, Sherafat Y, Peng B, Li X, Choi JH, Gou L, Zingg B, Azam S, Lo D, Khanjani N, Zhang B, Stanis J, Bowman I, Cotter K, Cao C, Yamashita S, Tugangui A, Li A, Jiang T, Jia X, Feng Z, Aquino S, Mun HS, Zhu M, Santarelli A, Benavidez NL, Song M, Dan G, Fayzullina M, Ustrell S, Boesen T, Johnson DL, Xu H, Bienkowski MS, Yang XW, Gong H, Levine MS, Wickersham I, Luo Q, Hahn JD, Lim BK, Zhang LI, Cepeda C, Hintiryan H, Dong HW (2021) The mouse cortico-basal ganglia-thalamic network. Nature 598:188–194. DOI:10.1038/s41586-021-03993-3.

Franklin KBJ, Paxinos G (2008) The mouse brain in stereotaxic coordinates, 3rd Edition. New York, NY, USA: Elsevier Academic Press.

Frégnac Y (2017) Big data and the industrialization of neuroscience: A safe roadmap for understanding the brain? Science 358:470–477. DOI:10.1126/science.aan8866.

Friston KJ, Frith CD, Passingham RE, Liddle PF, Frackowiak RS (1992) Motor practice and neurophysiological adaptation in the cerebellum: a positron tomography study. Proc Biol Sci 248:223–228. DOI:10.1098/rspb.1992.0065.

Garifoli A, Cardile V, Maci T, Perciavalle V (2003) Exercise increases cytochrome oxidase activity in specific cerebellar areas of the rat. Arch Ital Biol 141:181–187.

Gomes-Osman J, Cabral DF, Morris TP, McInerney K, Cahalin LP, Rundek T, Oliveira A, Pascual-Leone A (2018) Exercise for cognitive brain health in aging: A systematic review for an evaluation of dose. Neurol Clin Pract 8:257–265. DOI:10.1212/CPJ.0000000000000460.

Gopinath K, Krishnamurthy V, Cabanban R, Crosson BA (2015) Hubs of Anticorrelation in High-Resolution Resting-State Functional Connectivity Network Architecture. Brain Connect 5:267–275. DOI:10.1089/brain.2014.0323.

Grospe GM, Baker PM, Ragozzino ME (2018) Cognitive Flexibility Deficits Following 6-OHDA Lesions of the Rat Dorsomedial Striatum. Neuroscience 374:80–90. DOI:10.1016/j.neuroscience.2018.01.032.

Guo Y, Wang Z, Prathap S, Holschneider DP (2017) Recruitment of prefrontal-striatal circuit in response to skilled motor challenge. Neuroreport 28:1187–1194. DOI:10.1097/WNR.0000000000000881.

Hillman CH, Erickson KI, Kramer AF (2008) Be smart, exercise your heart: exercise effects on brain and cognition. Nat Rev Neurosci 9:58–65. DOI:10.1038/nrn2298.

Hintiryan H, Foster NN, Bowman I, Bay M, Song MY, Gou L, Yamashita S, Bienkowski MS, Zingg B, Zhu M, Yang XW, Shih JC, Toga AW, Dong HW (2016) The mouse cortico-striatal projectome. Nat Neurosci 19:1100–1114. DOI:10.1038/nn.4332.

Holman SR, Staines WR (2021) The effect of acute aerobic exercise on the consolidation of motor memories. Exp Brain Res 239:2461–2475. DOI:10.1007/s00221-021-06148-y.

Holschneider DP, Wang Z, Pang RD (2014) Functional connectivity-based parcellation and connectome of cortical midline structures in the mouse: a perfusion autoradiography study. Front Neuroinform 8:61. DOI:10.3389/fninf.2014.00061.

Holschneider DP, Yang J, Guo Y, Maarek JM (2007) Reorganization of functional brain maps after exercise training: Importance of cerebellar-thalamic-cortical pathway. Brain Res 1184:96–107. DOI:10.1016/j.brainres.2007.09.081.

Holschneider DP, Guo Y, Wang Z, Vidal M, Scremin OU (2019) Positive Allosteric Modulation of Cholinergic Receptors Improves Spatial Learning after Cortical Contusion Injury in Mice. J Neurotrauma 36:2233–2245. DOI:10.1089/neu.2018.6036.

Honey CJ, Kotter R, Breakspear M, Sporns O (2007) Network structure of cerebral cortex shapes functional connectivity on multiple time scales. Proc Natl Acad Sci U S A, 2007. 104(24):10240–5. doi: 10.1073/pnas.0701519104.

Hunnicutt BJ, Jongbloets BC, Birdsong WT, Gertz KJ, Zhong H, Mao T (2016) A comprehensive excitatory input map of the striatum reveals novel functional organization. Elife 5. DOI:10.7554/eLife.19103.

Hutchison RM, Womelsdorf T, Allen EA, Bandettini PA, Calhoun VD, Corbetta M, Della Penna S, Duyn JH, Glover GH, Gonzalez-Castillo J, Handwerker DA, Keilholz S, Kiviniemi V, Leopold DA, de Pasquale F, Sporns O, Walter M, Chang C (2013) Dynamic functional connectivity: promise, issues, and interpretations. Neuroimage 80:360–378. DOI:10.1016/j.neuroimage.2013.05.079.

Inoue T, Ninuma S, Hayashi M, Okuda A, Asaka T, Maejima H (2018) Effects of long-term exercise and low-level inhibition of GABAergic synapses on motor control and the expression of BDNF in the motor related cortex. Neurol Res 40:18–25. DOI:10.1080/01616412.2017.1382801.

Jancke L, Shah NJ, Peters M (2000) Cortical activations in primary and secondary motor areas for complex bimanual movements in professional pianists. Brain Res Cogn Brain Res 10:177–183. DOI:10.1016/s0926-6410(00)00028-8.

Jenkins IH, Brooks DJ, Nixon PD, Frackowiak RS, Passingham RE (1994) Motor sequence learning: a study with positron emission tomography. J Neurosci 14:3775–3790. DOI:10.1523/JNEUROSCI.14-06-03775.1994.

Ji L, Zhang H, Potter GG, Zang YF, Steffens DC, Guo H, Wang L (2017) Multiple Neuroimaging Measures for Examining Exercise-induced Neuroplasticity in Older Adults: A Quasi-experimental Study. Front Aging Neurosci 9:102. DOI:10.3389/fnagi.2017.00102.

Karni A, Meyer G, Rey-Hipolito C, Jezzard P, Adams MM, Turner R, Ungerleider LG (1998) The acquisition of skilled motor performance: fast and slow experience-driven changes in primary motor cortex. Proc Natl Acad Sci U S A 95:861–868. DOI:10.1073/pnas.95.3.861.

Kim DE, Shin MJ, Lee KM, Chu K, Woo SH, Kim YR, Song EC, Lee JW, Park SH, Roh JK (2004) Musical training-induced functional reorganization of the adult brain: functional magnetic resonance imaging and transcranial magnetic stimulation study on amateur string players. Hum Brain Mapp 23:188–199. DOI:10.1002/hbm.20058.

Knox JE, Harris KD, Graddis N, Whitesell JD, Zeng H, Harris JA, Shea-Brown E, Mihalas S (2019) High-resolution data-driven model of the mouse connectome. Netw Neurosci 3:217–236. DOI:10.1162/netn_a_00066.

Koeneke S, Lutz K, Wustenberg T, Jancke L (2004) Long-term training affects cerebellar processing in skilled keyboard players. Neuroreport 15:1279–1282. DOI:10.1097/01.wnr.0000127463.10147.e7.

Lehmann N, Villringer A, Taubert M (2020) Colocalized White Matter Plasticity and Increased Cerebral Blood Flow Mediate the Beneficial Effect of Cardiovascular Exercise on Long-Term Motor Learning. J Neurosci 40:2416–2429. DOI:10.1523/JNEUROSCI.2310-19.2020.

Li HQ, Spitzer NC (2020) Exercise enhances motor skill learning by neurotransmitter switching in the adult midbrain. Nat Commun 11:2195. DOI:10.1038/s41467-020-16053-7.

Liang Z, King J, Zhang N (2012) Anticorrelated resting-state functional connectivity in awake rat brain. Neuroimage 59:1190–1199. DOI:10.1016/j.neuroimage.2011.08.009.

Loras H, Haga M, Sigmundsson H (2020) Effect of a Single Bout of Acute Aerobic Exercise at Moderate-to-Vigorous Intensities on Motor Learning, Retention and Transfer. Sports (Basel) 8. DOI:10.3390/sports8020015.

Ludyga S, Gerber M, Puhse U, Looser VN, Kamijo K (2020) Systematic review and meta-analysis investigating moderators of long-term effects of exercise on cognition in healthy individuals. Nat Hum Behav 4:603–612. DOI:10.1038/s41562-020-0851-8.

Lundquist AJ, Parizher J, Petzinger GM, Jakowec MW (2019) Exercise induces region-specific remodeling of astrocyte morphology and reactive astrocyte gene expression patterns in male mice. J Neurosci Res 97:1081–1094. DOI:10.1002/jnr.24430.

Ma L, Wang B, Narayana S, Hazeltine E, Chen X, Robin DA, Fox PT, Xiong J (2010) Changes in regional activity are accompanied with changes in inter-regional connectivity during 4 weeks motor learning. Brain Res 1318:64–76. DOI:10.1016/j.brainres.2009.12.073.

Magon S, Donath L, Gaetano L, Thoeni A, Radue EW, Faude O, Sprenger T (2016) Striatal functional connectivity changes following specific balance training in elderly people: MRI results of a randomized controlled pilot study. Gait Posture 49:334–339. DOI:10.1016/j.gaitpost.2016.07.016.

McCloskey DP, Adamo DS, Anderson BJ (2001) Exercise increases metabolic capacity in the motor cortex and striatum, but not in the hippocampus. Brain Res 891:168–175. DOI:10.1016/s0006-8993(00)03200-5.

McNamara A, Tegenthoff M, Dinse H, Buchel C, Binkofski F, Ragert P (2007) Increased functional connectivity is crucial for learning novel muscle synergies. Neuroimage 35:1211–1218. DOI:10.1016/j.neuroimage.2007.01.009.

Mills BD, Grayson DS, Shunmugavel A, Miranda-Dominguez O, Feczko E, Earl E, Neve KA, Fair DA (2018) Correlated Gene Expression and Anatomical Communication Support Synchronized Brain Activity in the Mouse Functional Connectome. J Neurosci 38:5774–5787. DOI:10.1523/JNEUROSCI.2910-17.2018.

Moore D, Jung M, Hillman CH, Kang M, Loprinzi PD (2022) Interrelationships between exercise, functional connectivity, and cognition among healthy adults: A systematic review. Psychophysiology 59:e14014. DOI:10.1111/psyp.14014.

Morgen K, Kadom N, Sawaki L, Tessitore A, Ohayon J, Frank J, McFarland H, Martin R, Cohen LG (2004) Kinematic specificity of cortical reorganization associated with motor training. Neuroimage 21:1182–1187. DOI:10.1016/j.neuroimage.2003.11.006.

Munte TF, Altenmuller E, Jancke L (2002) The musician’s brain as a model of neuroplasticity. Nat Rev Neurosci 3:473–478. DOI:10.1038/nrn843.

Nair HP, Gonzalez-Lima F (1999) Extinction of behavior in infant rats: development of functional coupling between septal, hippocampal, and ventral tegmental regions. J Neurosci 19:8646–8655. DOI:10.1523/JNEUROSCI.19-19-08646.1999.

Navarro A, Gomez C, Lopez-Cepero JM, Boveris A (2004) Beneficial effects of moderate exercise on mice aging: survival, behavior, oxidative stress, and mitochondrial electron transfer. Am J Physiol Regul Integr Comp Physiol 286:R505–511. DOI:10.1152/ajpregu.00208.2003.

Needham BD, Funabashi M, Adame MD, Wang Z, Boktor JC, Haney J, Wu WL, Rabut C, Ladinsky MS, Hwang SJ, Guo Y, Zhu Q, Griffiths JA, Knight R, Bjorkman PJ, Shapiro MG, Geschwind DH, Holschneider DP, Fischbach MA, Mazmanian SK (2022) A gut-derived metabolite alters brain activity and anxiety behaviour in mice. Nature 602:647–653. DOI:10.1038/s41586-022-04396-8.

Nguyen PT, Holschneider DP, Maarek JM, Yang J, Mandelkern MA (2004) Statistical parametric mapping applied to an autoradiographic study of cerebral activation during treadmill walking in rats. Neuroimage 23:252–259. DOI:10.1016/j.neuroimage.2004.05.014.

Nicolini C, Fahnestock M, Gibala MJ, Nelson AJ (2021) Understanding the Neurophysiological and Molecular Mechanisms of Exercise-Induced Neuroplasticity in Cortical and Descending Motor Pathways: Where Do We Stand? Neuroscience 457:259–282. DOI:10.1016/j.neuroscience.2020.12.013.

O’Neill M, Brown VJ (2007) The effect of striatal dopamine depletion and the adenosine A2A antagonist KW-6002 on reversal learning in rats. Neurobiol Learn Mem 88:75–81. DOI:10.1016/j.nlm.2007.03.003.

Oh SW, Harris JA, Ng L, Winslow B, Cain N, Mihalas S, Wang Q, Lau C, Kuan L, Henry AM, Mortrud MT, Ouellette B, Nguyen TN, Sorensen SA, Slaughterbeck CR, Wakeman W, Li Y, Feng D, Ho A, Nicholas E, Hirokawa KE, Bohn P, Joines KM, Peng H, Hawrylycz MJ, Phillips JW, Hohmann JG, Wohnoutka P, Gerfen CR, Koch C, Bernard A, Dang C, Jones AR, Zeng H (2014) A mesoscale connectome of the mouse brain. Nature 508:207–214. DOI:10.1038/nature13186.

Peng YH, Heintz R, Wang Z, Guo Y, Myers KG, Scremin OU, Maarek JM, Holschneider DP (2014) Exercise training reinstates cortico-cortical sensorimotor functional connectivity following striatal lesioning: development and application of a subregional-level analytic toolbox for perfusion autoradiographs of the rat brain. Front Phys 2. DOI:10.3389/fphy.2014.00072.

Petzinger GM, Fisher BE, McEwen S, Beeler JA, Walsh JP, Jakowec MW (2013) Exercise-enhanced neuroplasticity targeting motor and cognitive circuitry in Parkinson’s disease. Lancet Neurol 12:716–726. DOI:10.1016/S1474-4422(13)70123-6.

Pinho AL, de Manzano O, Fransson P, Eriksson H, Ullen F (2014) Connecting to create: expertise in musical improvisation is associated with increased functional connectivity between premotor and prefrontal areas. J Neurosci 34:6156–6163. DOI:10.1523/JNEUROSCI.4769-13.2014.

Puga F, Barrett DW, Bastida CC, Gonzalez-Lima F (2007) Functional networks underlying latent inhibition learning in the mouse brain. Neuroimage 38:171–183. DOI:10.1016/j.neuroimage.2007.06.031.

Quaney BM, Boyd LA, McDowd JM, Zahner LH, He J, Mayo MS, Macko RF (2009) Aerobic exercise improves cognition and motor function poststroke. Neurorehabil Neural Repair 23:879–885. DOI:10.1177/1545968309338193.

Schwarz AJ, Gozzi A, Reese T, Heidbreder CA, Bifone A (2007) Pharmacological modulation of functional connectivity: the correlation structure underlying the phMRI response to d-amphetamine modified by selective dopamine D3 receptor antagonist SB277011A. Magn Reson Imaging 25:811–820. DOI:10.1016/j.mri.2007.02.017.

Seidler RD (2004) Multiple motor learning experiences enhance motor adaptability. J Cogn Neurosci 16:65–73. DOI:10.1162/089892904322755566.

Shumake J, Conejo-Jimenez N, Gonzalez-Pardo H, Gonzalez-Lima F (2004) Brain differences in newborn rats predisposed to helpless and depressive behavior. Brain Res 1030:267–276. DOI:10.1016/j.brainres.2004.10.015.

Sokoloff L (1991) Measurement of local cerebral glucose utilization and its relation to local functional activity in the brain. Adv Exp Med Biol 291:21–42. DOI:10.1007/978-1-4684-5931-9_4.

Sokoloff L, Reivich M, Kennedy C, Des Rosiers MH, Patlak CS, Pettigrew KD, Sakurada O, Shinohara M (1977) The [14C]deoxyglucose method for the measurement of local cerebral glucose utilization: theory, procedure, and normal values in the conscious and anesthetized albino rat. J Neurochem 28:897–916. DOI:10.1111/j.1471-4159.1977.tb10649.x.

Soncrant TT, Horwitz B, Holloway HW, Rapoport SI (1986) The pattern of functional coupling of brain regions in the awake rat. Brain Res 369:1–11. DOI:10.1016/0006-8993(86)90507-x.

Stafford JM, Jarrett BR, Miranda-Dominguez O, Mills BD, Cain N, Mihalas S, Lahvis GP, Lattal KM, Mitchell SH, David SV, Fryer JD, Nigg JT, Fair DA (2014) Large-scale topology and the default mode network in the mouse connectome. Proc Natl Acad Sci U S A 111:18745–18750. DOI:10.1073/pnas.1404346111.

Sun FT, Miller LM, Rao AA, D’Esposito M (2007) Functional connectivity of cortical networks involved in bimanual motor sequence learning. Cereb Cortex 17:1227–1234. DOI:10.1093/cercor/bhl033.

Tao J, Chen X, Egorova N, Liu J, Xue X, Wang Q, Zheng G, Li M, Hong W, Sun S, Chen L, Kong J (2017) Tai Chi Chuan and Baduanjin practice modulates functional connectivity of the cognitive control network in older adults. Sci Rep 7:41581. DOI:10.1038/srep41581.

Tracy JI, Faro SS, Mohammed F, Pinus A, Christensen H, Burkland D (2001) A comparison of ‘Early’ and ‘Late’ stage brain activation during brief practice of a simple motor task. Brain Res Cogn Brain Res 10:303–316. DOI:10.1016/s0926-6410(00)00045-8.

Trujillo JP, Gerrits NJ, Veltman DJ, Berendse HW, van der Werf YD, van den Heuvel OA (2015) Reduced neural connectivity but increased task-related activity during working memory in de novo Parkinson patients. Hum Brain Mapp 36:1554–1566. DOI:10.1002/hbm.22723.

Venkadesh S, Van Horn JD (2021) Integrative Models of Brain Structure and Dynamics: Concepts, Challenges, and Methods. Front Neurosci 15:752332. DOI:10.3389/fnins.2021.752332.

Vissing J, Andersen M, Diemer NH (1996) Exercise-induced changes in local cerebral glucose utilization in the rat. J Cereb Blood Flow Metab 16:729–736. DOI:10.1097/00004647-199607000-00025.

Wang Z, Pang RD, Hernandez M, Ocampo MA, Holschneider DP (2012) Anxiolytic-like effect of pregabalin on unconditioned fear in the rat: an autoradiographic brain perfusion mapping and functional connectivity study. Neuroimage 59:4168–4188. DOI:10.1016/j.neuroimage.2011.11.047.

Wang Z, Bradesi S, Charles JR, Pang RD, Maarek JI, Mayer EA, Holschneider DP (2011) Functional brain activation during retrieval of visceral pain-conditioned passive avoidance in the rat. Pain 152:2746–2756. DOI:10.1016/j.pain.2011.08.022.

Wang Z, Guo Y, Myers KG, Heintz R, Peng YH, Maarek JM, Holschneider DP (2015) Exercise alters resting-state functional connectivity of motor circuits in parkinsonian rats. Neurobiol Aging 36:536–544. DOI:10.1016/j.neurobiolaging.2014.08.016.

Wehrl HF, Hossain M, Lankes K, Liu CC, Bezrukov I, Martirosian P, Schick F, Reischl G, Pichler BJ (2013) Simultaneous PET-MRI reveals brain function in activated and resting state on metabolic, hemodynamic and multiple temporal scales. Nat Med 19:1184–1189. DOI:10.1038/nm.3290.

Wheeler AL, Teixeira CM, Wang AH, Xiong X, Kovacevic N, Lerch JP, McIntosh AR, Parkinson J, Frankland PW (2013) Identification of a functional connectome for long-term fear memory in mice. PLoS Comput Biol 9:e1002853. DOI:10.1371/journal.pcbi.1002853.

Won J, Callow DD, Pena GS, Gogniat MA, Kommula Y, Arnold-Nedimala NA, Jordan LS, Smith JC (2021) Evidence for exercise-related plasticity in functional and structural neural network connectivity. Neurosci Biobehav Rev 131:923–940. DOI:10.1016/j.neubiorev.2021.10.013.

Yang Z, Zhu T, Pompilus M, Fu Y, Zhu J, Arjona K, Arja RD, Grudny MM, Plant HD, Bose P, Wang KK, Febo M (2021) Compensatory functional connectome changes in a rat model of traumatic brain injury. Brain Commun 3:fcab244. DOI:10.1093/braincomms/fcab244.

Zerbi V, Floriou-Servou A, Markicevic M, Vermeiren Y, Sturman O, Privitera M, von Ziegler L, Ferrari KD, Weber B, De Deyn PP, Wenderoth N, Bohacek J (2019) Rapid Reconfiguration of the Functional Connectome after Chemogenetic Locus Coeruleus Activation. Neuron 103:702–718 e705. DOI:10.1016/j.neuron.2019.05.034.

Zingg B, Hintiryan H, Gou L, Song MY, Bay M, Bienkowski MS, Foster NN, Yamashita S, Bowman I, Toga AW, Dong HW (2014) Neural networks of the mouse neocortex. Cell 156:1096–1111. DOI:10.1016/j.cell.2014.02.023.

