## Supplemental Table S1 for "Exercise alters cortico-basal ganglia network functional connectivity: A mesoscopic level analysis informed by anatomic parcellation defined in the mouse brain connectome"

**Supplemental Table S1. Intra-structural functional connectivity density.** Connectivity density is defined as the number of connections expressed as a percentage of the total number of possible connections within that structure.

|  |  | **Caudo-putamen** | **Substantia nigra pars reticulata** | **Globus pallidus externus** | **Motor cortex** | **Prefrontal cortex** | **Thalamus** |
| --- | --- | --- | --- | --- | --- | --- | --- |
| **Positive (%)** | Exercise  Control | +20.0  +11.0 | +51.7 +45.7 | +51.6 +58.1 | +29.4 +23.5 | +31.6 +19.3 | +22.4 +48.6 |
| **Negative (%)** | Exercise  Control | -0.1  -2.1 | -0.0  -0.0 | -0.0  -0.0 | -0.0  -2.0 | -0.0  -1.2 | -0.0  -0.0 |
