## Supplemental Table S2 for "Exercise alters cortico-basal ganglia network functional connectivity: A mesoscopic level analysis informed by anatomic parcellation defined in the mouse brain connectome"

**Supplemental Table S2. Inter-structural functional connectivity density between specific structures.**

The top-right half of the table shows positive connectivity densities, which are marked with "+". The lower-left half of the table shows negative connectivity densities, marked with "-". Connectivity densities are expressed as a percentage of the total number of possible connections between two specified brain structures.

|  |  |  | **CP** | **SNr** | **GPe** | **M1/M2** | **PFC** | **Thal** |
| --- | --- | --- | --- | --- | --- | --- | --- | --- |
| **Caudoputamen** | Exercise  Control |  |  | +0.0  +4.4 | +12.8  +13.9 | +14.5  +0.8 | +3.6  +1.2 | +2.7  +1.7 |
| **Substantia nigra pars reticulata** | Exercise  Control |  | -6.3  -1.9 |  | +0.0  +3.0 | +0.7  +1.6 | +1.7  +1.3 | +0.4  +0.0 |
| **Globus pallidus externus** | Exercise  Control |  | -0.1  -0.2 | -18.5  -2.4 |  | +0.0  +1.9 | +0.0  +0.8 | +3.4  +0.7 |
| **Motor cortex** | Exercise  Control |  | -0.3  -0.4 | -0.9  -0.9 | -2.5  -1.0 |  | +5.0  +9.6 | +0.3  +12.7 |
| **Prefrontal cortex** | Exercise  Control |  | -0.6  -7.8 | -0.4  -7.4 | -0.0  -21.4 | -0.3  -2.0 |  | +0.3  +17.0 |
| **Thalamus** | Exercise  Control |  | -5.4  -4.7 | -0.0  -3.8 | -3.4  -3.9 | -5.3  -0.8 | -7.3  -0.0 |  |
